# Comparison of DNA targeting CRISPR editors in human cells

**DOI:** 10.1101/2022.10.20.513037

**Authors:** Hongxin Huang, Weiqi Lv, Jinhe Li, Guanjie Huang, Zhihong Tan, Yongfei Hu, Shufeng Ma, Xin Zhang, Linxuan Huang, Ying Lin

## Abstract

Profiling and comparing the performance of current widely used DNA targeting CRISPR systems provide the basic information for the gene-editing toolkit and can be a useful resource for this field. Herein we made a parallel comparison between the recently reported miniature Cas12f1 and the widely used Cas12a and Cas9 nucleases in mammalian cells. The results revealed that as a CRISPRa activator, Un1Cas12f1 could induce gene expression with a comparable level to these of Cas12a and Cas9, while as a cleaving editor, Cas12f1 exhibited similar properties to Cas12a, like high specificity and dominantly induced deletions over insertions, but with less activity. In contrast, wild-type SpCas9 showed the highest activity, lowest specificity, and induced balanced deletions over insertions. Thus, Cas12f1 is recommended for gene-activation-based applications, Cas12a is for therapy applications, and wild-type Cas9 is for *in vitro* and animal investigations. The comparison provided the editing properties of current widely used DNA-targeting CRISPR systems for the gene-editing field.

## Introduction

DNA-targeting CRISPR systems have been developed as powerful tools for basic research and clinical therapy, including programmable DNA editing, gene activation/suppression, live imaging, base editing, and primer editing^1, 2^. However, therapeutic delivery of these systems remains challenging, in part because their sizes exceed the packaging capacity (<4.7 kb) of adeno-associated virus (AAV), the most widely used viral vector for gene delivery. To overcome this limitation, efforts have been made to explore the miniature Cas-nucleases, such as the *Sa*Cas9 (1053 amino acids)^3^, *Cj*Cas9 (984 amino acids)^4^, and Cas12j (700-800 amino acids)^5^, et al.. However, the low editing activity or the relatively large size of these nucleases keep the challenge incompletely solved.

Recently, it has been reported that the type V-F Cas12f (also known as Cas14) nuclease could serve as a hypercompact gene-editing tool in mammalian cells with optimized sgRNA or engineered nuclease mutant^6–10^. Cas12f forms a unique asymmetric dimer structure to bind crRNA and guide target DNA recognition and cleavage^11, 12^. The extremely small size (422 and 529 amino acids for *As*Cas12f1 and Un1Cas12f1, respectively, the two nucleases functional in mammalian cells) makes Cas12f1 an ideal tool for therapeutic editing. However, the editing features of these miniature nucleases in mammalian or human cells have not been well elucidated, such as the editing activity, specificity, cutting site, indel property, and transcriptional activation ability, leaving the question that how to choose them appropriately for a specific application.

In the current study, we analyzed the editing features of Cas12f1 and made a comparison between Cas12f1 and the other two widely used nucleases, Cas9 and Cas12a. Our data profiled the properties of current widely used DNA targeting CRISPR systems, which can be a guideline and a useful resource for the gene-editing field.

## Results and discussion

### Profiling the cutting features of the engineered CRISPR-Un1Cas12f1 systems

First, we used the WT-Un1Cas12f1 (G297C) Cas-nuclease and its mutant version V3.1 (D143R/T147R/E151A/G297C), both of which were from a previous report^8^, to combine with three engineered sgRNAs, ge3.0, ge4.0, and ge4.1 (Fig. 1a and Supplementary Fig. 1a-d)^9^, since these engineered nucleases had been proven to boost the genome regulation and editing ability of the CRISPR-Cas12f1 effector^8^, and the three modified sgRNAs were reported to improve for robust genome-editing^9^, however, the performance of their combinations had not been defined. The Tag-seq approach^13^, which detects the Cas-induced double-strand breaks (DSBs) by simply tracing the integrated exogenous oligonucleotide (termed Tag) via next-generation-sequencing, is a convenient and scalable method for genome-wide specificity assessment of CRISPR editors, and can accurately identify and characterize Cas-induced DSBs (Supplementary Fig. 1e). Thus, after confirming the comparable Un1Cas12f1-protein levels by western blotting (Supplementary Fig. 2), we employed this method to examine the editing abilities of the engineered Un1Cas12f1 combinations in HEK293T cells with twenty-one sgRNAs targeting eighteen genes (Fig. 1b and Supplementary Fig. 3, 4). Because most of these sites were well-tested in the previous studies^8, 9^. Since Tag-seq can accurately profile the Cas-induced DSBs, we first detected this property of the engineered Un1Cas12f1 editors. Consistent with the previous *in vitro* assays^7^, Tag-seq showed that the integrated Tag DNA oligonucleotides were multiple staggered breaks and were distal to the TTTR (R=A/T) PAM sequence, approximately peaking at the region of 14^th^ bp and 25^th^ bp downstream of the TTTR PAM (Fig. 1c and Supplementary Fig. 5), indicating the break sites induced by Cas12f1 nucleases were at this region. Additionally, the data also revealed that the region around the 8^th^ bp could be a new cut site for *in vivo* editing (Fig. 1c), suggesting that there may be a new cutting pattern for the Un1Cas12f1 editor, which was a little distinct from its *in vitro* assays^7^.

**Figure 1.**
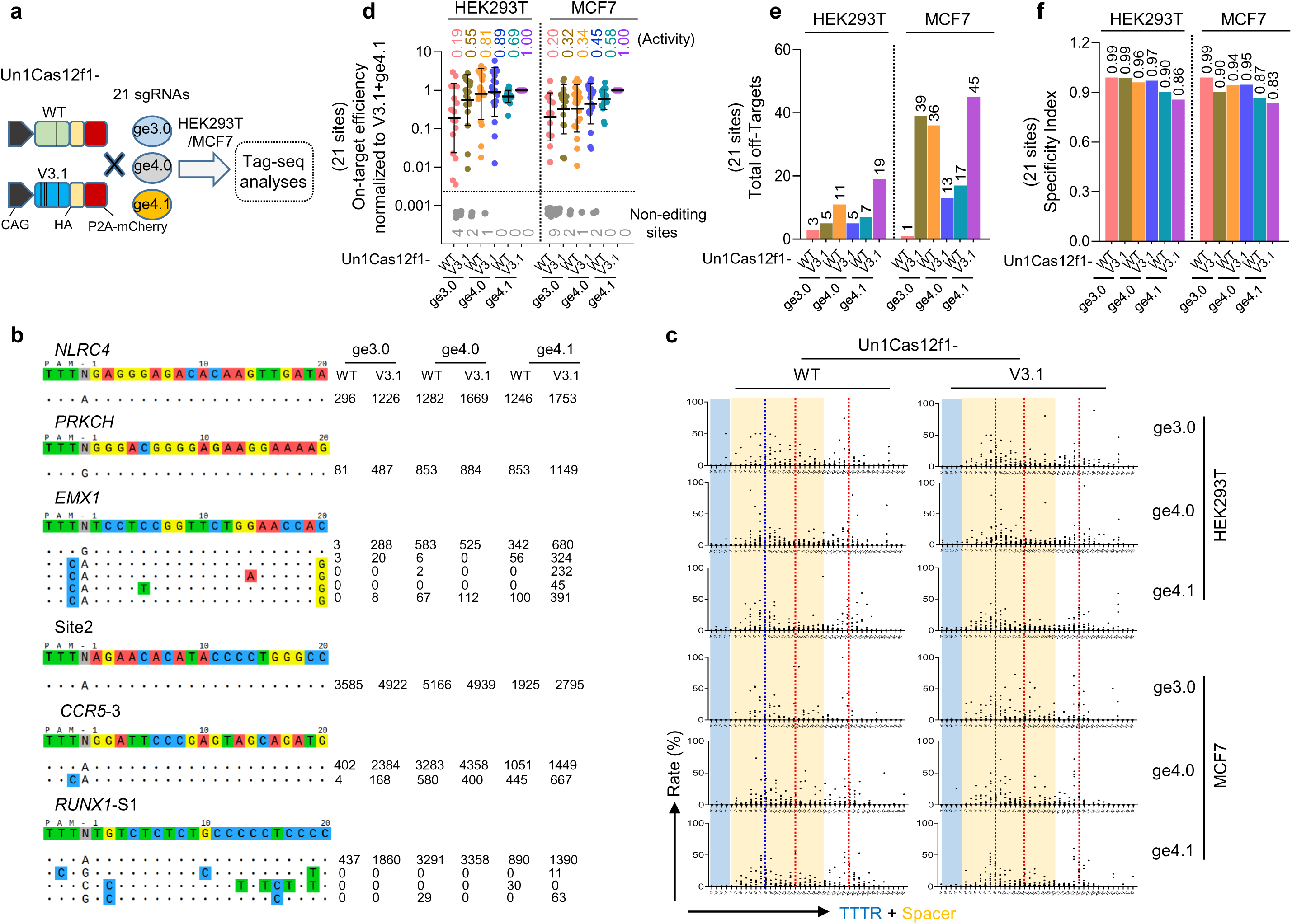
The performance of the CRISPR-Un1Cas12f1 systems in human cells. **a** Schematic of the editing ability analysis of the engineered CRISPR-Un1Cas12f1 systems, which contain two nucleases (also see Supplementary Fig. 1a) combined with three engineered sgRNA, ge3.0, ge4.0, and ge4.1. **b** Tag-seq-based comparative analysis of wild-type Un1Cas12f1 (WT), and Un1Cas12f1-V3.1 (V3.1, a reported high-active variant) combined with the engineered sgRNAs (ge3.0, ge4.0, and ge4.1) targeted to twenty-one sites in HEK293T cells (also see Supplementary Fig. 4). The sgRNA reference is shown on the top and the on-target and the off-target cleavages are displayed without or with mismatches to the sgRNA reference by color highlighting. Sequencing read counts are shown to the right of each site. **c** The characteristics of the Un1Cas12f1 induced double-strand-breaks (DSBs). Each point was calculated as the ratio of the read count at each break site to the total break read counts at this sgRNA, and then pooled within the twenty-one sgRNAs. Read counts were obtained from Tag-seq (Supplementary Fig. 5). x-axis shows the location of the sgRNA. The red dotted line indicates the reported *in vitro* DSB sites, while the blue dotted line indicates another potential breakpoint *in vivo*. **d** Normalization of on-target activity of the various CRISPR-Un1Cas12f1 systems to Un1Cas12f1-V3.1+ge4.1 combination, value= (other system on-target reads)/(V3.1+ge4.1 on-target reads). Grey points, sites without detectable editing. **e** Total number of off-target sites detected within the twenty-one sgRNAs. **f** Specificity Index assessment (value was calculated by the ratio of total on-target reads to the on-target reads plus the off-target reads within the twenty-one sites).

### Profiling the editing performance of the engineered CRISPR-Un1Cas12f1 systems

We then assessed the editing performance of these Un1Cas12f1 systems by Tag-seq results. Globally, the variant V3.1 Un1Cas12f1 displayed higher activity than that of WT Un1Cas12f1, and the engineered sgRNAs ge4.1 and ge4.0 were better than that of ge3.0 (Fig. 1d), both of which were consistent with the previous reports^8, 9^. In detail, the V3.1 combined with the ge4.1 sgRNA (V3.1+ge4.1) displayed a robust editing ability, since it can edit all the twenty-one tested sites and was the most active combination, the V3.1 combined with the ge4.0 was the second good (V3.1+ge4.0), and the contained ge3.0 sgRNA editors with several non-editing sites were the lowest one (Fig. 1d). For the specificity, although the ge3.0 versions exhibited the highest specificity index, which mainly resulted from its low activity (Fig. 1d-f). Generally, in line with the “trade-off” hypothesis that increased activity compromises specificity and *vice versa*^14, 15^, the V3.1+ge4.1 editor displayed the best efficiency but with the lowest specificity, while the V3.1+ge4.0 combination retained balanced in activity and specificity (Fig. 1d-f and Supplementary Fig. 4). Together, these data demonstrated that using a high-efficient Cas-nuclease variant combined with a better-engineered sgRNA could improve the performance of the Un1Cas12f1 system without badly damaging specificity. And similar results could be observed in another cell line MCF7 (Fig. 1d-f and Supplementary Fig. 5, 6). Thus, the V3.1+ge4.1 and V3.1+ge4.0 systems were selected for further study.

### Transcriptional activation with the DNA-targeting CRISPR editors

DNA-targeting CRISPR systems had been harnessed as versatile approaches for a variety of applications^1^, however, these systems’ editing features, such as activity, specificity, cutting site, indel property, and transcriptional activation ability, have not been parallelly and comprehensively elucidated. Thus, we next focused on several common DNA-targeting CRISPR systems, including CRISPR-*Sp*Cas9^16^, CRISPR-Cas12a (*As*Cas12a and *Lb*Cas12a)^17^, CRISPR-Un1Cas12f1 (V3.1+ge4.1 and V3.1+ge4.0)^8, 9^, and CRISPR-*As*Cas12f1 (another Cas12f1 editor)^7, 10, 18, 19^, to compare their gene-editing performance.

Since the CRISPR-based activation (CRISPRa) system is a useful technique, which holds great promise for clinical therapy applications ^20–22^, and the Cas12f1 system had been reported to be engineered as such an activator for robust gene activation^8^. Thus we first intended to determine the transcriptional activation abilities of these six CRISPRa activators. We constructed these DNA-targeting CRISPRa by fusing the DNase-inactive Cas-nuclease to the synthetic VPR (VP64-p65-Rta) activation domain and then tested their transcriptional activation of *IL1RN* and *HBG* in human cells HEK293T and MCF7, and *Fgf21* in mouse B16 cell line (Fig. 2a). As a result, we found that similar to the Cas12a and Cas9 CRISPRa systems, dUn1Cas12f-V3.1 combined with the ge4.1 or the ge4.0 sgRNA could induce *IL1RN*, *HBG, and Fgf21* expression with a comparable level in all the tested cells (Fig. 2b-d), which was consistent with the previous study^8^, demonstrating its ability in epigenetic regulation. Nevertheless, the dAsCas12f1-VPR system, which exhibited undetectable activity (Fig. 2b-d), implies its low capacity for gene regulation, and thus further engineering for improvement is required. These data suggested that the miniature CRISPR-Un1Cas12f1 system was also a powerful CRISPRa platform, which can be an alternative tool for gene activation.

**Figure 2.**
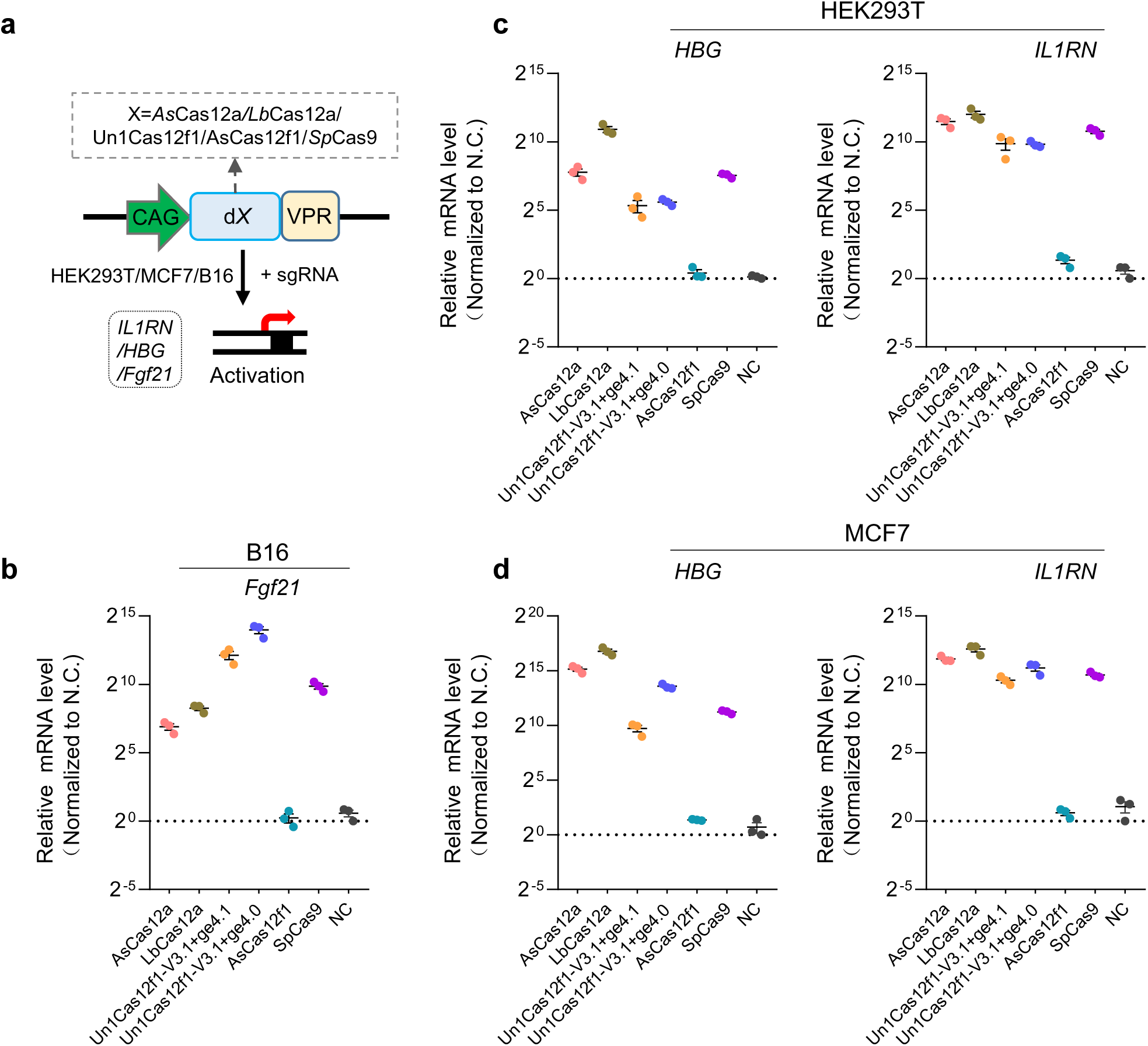
Gene activation comparison of the DNA targeting CRISPRa systems in mammalian cells. **a** Schematic of the gene activation system based on DNA targeting CRISPR systems. VPR, synthetic VP64-p65-Rta activation domain. **b-d** qPCR analyses of the transcriptional activation levels with DNA targeting CRISPRa guided by a single sgRNA targeting each promoter region of *Fgf21* mouse B16 cells (b), and *IL1RN* and *HBG* in human cells HEK293T (c) and MCF7 (d), respectively. Mean values are presented with SEM, n=3 independent experiments.

### Activities comparison with the DNA-targeting CRISPR editors

Next, we intended to examine the editing activities of these six DNA editors, because the editing efficiency was a key concern for a CRISPR-Cas gene-editing tool. After confirming the similar Cas-protein expression level and transfection efficiency (Supplementary Fig. 2, 3), the comparison was performed by Deep-seq experiment in HEK293T cells (Fig. 3a). To enable a better comparison of the on-target activities of the miniature Cas12f1 with that of widely-used Cas9 and Cas12a, we designed twenty-one endogenous human gene targeting sites that contained overlapping spacers for both Cas9 and Cas12a/Cas12f1 nucleases, where the sequences started with the TTTR motif (locations should be at -1 to -4) and ended with the NGG PAM (locations should be at 20 to 23 bp). And these sites were chosen to have variable numbers of predicted off-target sites in the genome, as estimated using Cas-OFFinder^23^ (Table S1). Consistent with the recently reported study^24^, Deep-seq results exhibited that the *Sp*Cas9 enzyme was the most active in all the tested Cas-nucleases, followed by Cas12a, and then Cas12f1 (Fig. 3b and Supplementary Fig. 7). However, for the Un1Cas12f1 editors, although the best Cas12f1/gRNA combination was used, its activity still displayed limited, which was much lower than that of *Sp*Cas9 and Cas12a (Fig. 3b), and the disruption of mNeonGreen expression in a HEK293T knock-in reporter cell line also confirmed the poor editing activity of Cas12f1 (Fig. 3c and Supplementary Fig. 8), both of which were quite different from the previous and the recently reported investigations. We speculated the most possible reason might come from the transfection method. In the recently reported work, they determined the editing efficiency within 2-7 days after 3 days of puromycin selection by single sgRNA transfection (thus the transfection efficiency = ∼100%), while we calculated the cutting activity 2 days post-transfection by employing a scalable method that the experiment was administrated by pooling all the twenty-one sgRNAs with a single transfection (the total amount of the input DNA was the same as a single guide and the transfection efficiency = ∼30-70%, Supplementary Fig. 3). Because parallelly measuring the editing efficiency at diverse sites in a single transfection was a rapid and cost-efficient approach for evaluating the performance of a nuclease^13, 14^. In addition, the Cas12f1 proteins used in these two works were a little different. In our investigation, we employed the Un1Cas12f1 nucleases, which contained a G297C mutation, were originally from another recent study on Un1Cas12f1^8^. In terms of inducing indels, Deep-seq data showed that *Sp*Cas9 was more likely to produce balanced insertions and deletions because it led to blunt end breaks^16, 25^, while Cas12f1 and Cas12a tended to induce deletions since they generated sticky end breaks (Fig. 3d and Supplementary Fig. 7) ^7–10, 17, 19, 26^. Exception for HEK293T cells, we also performed the experiments in other human cell lines, such as MCF7, K562, and Jurkat (Fig. 3d-f and Supplementary Fig. 7), and the same results were obtained.

**Figure 3.**
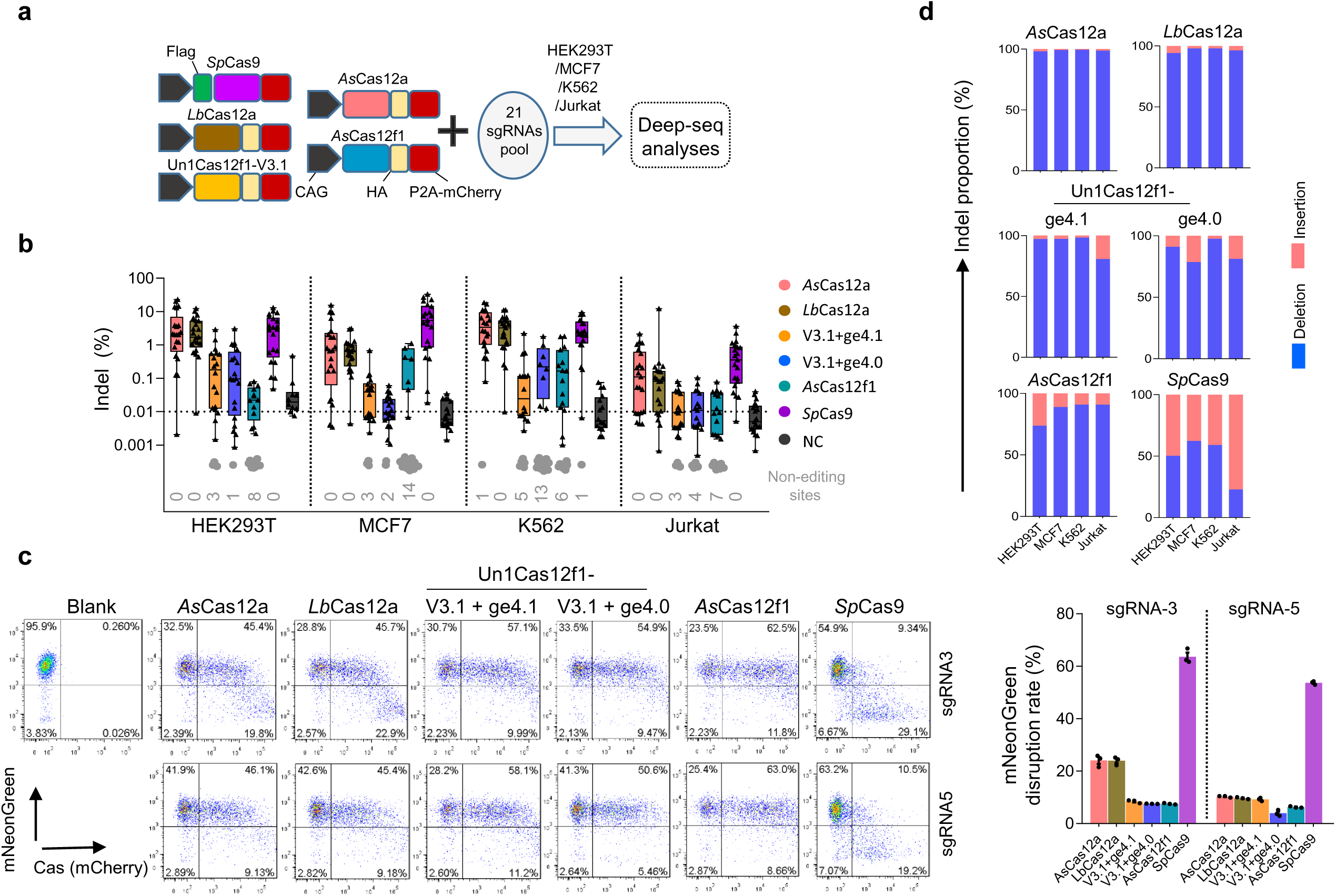
Editing activities comparison of the DNA targeting CRISPR systems in human cells. **a** Schematic of the editing performance analysis by Deep-seq among the DNA targeting CRISPR editors (CRISPR-*As*Cas12a, CRISPR-*Lb*Cas12a, CRISPR-Un1Cas12f1, CRISPR-*As*Cas12f1, and CRISPR-*Sp*Cas9 systems). **b** The editing activities of the DNA targeting systems with twenty-one sgRNAs in HEK293T, MCF7, K562, and Jurkat cells were revealed by Deep-seq (also see Supplementary Fig. 7). The targeted sgRNAs for Cas12 and Cas9 share a common spacer sequence. Grey points, sites without detectable editing. **c** FACS revealed the editing activities of the DNA targeting systems by disruption of mNeonGreen expression in a HEK293T-KI reporter cell line. The editing efficiency was determined as the proportion of GFP negative cells within the Cas-nucleases transfected cells (mCherry-positive). sgRNA3/5, sgRNAs targeting mNeonGreen. Mean values are presented with SEM, n=3 independent experiments (also see Supplementary Fig. 8, another two replicates). **d** The proportion of the insertions and deletions induced by the DNA targeting systems within the twenty-one sites.

### Specificities comparison among the DNA-targeting CRISPR editors

As the off-target effect is another major concern of the CRISPR-Cas systems for therapeutic applications, we finally wanted to know the targeting accuracy of these six editors. Thus we again performed Tag-seq assays with the above twenty-one sgRNAs to compare their specificities in genome-editing (Fig. 4a, b and Supplementary Fig. 3 and 9). As a result, for the cutting features, Tag-seq showed that the break site induced by *Sp*Cas9 was at 3^th^-4^th^ bp upstream of its NGG PAM, and that Cas12a (*As*Cas12a and *Lb*Cas12a) displayed multiple staggered breaks peaking at around 14^th^ and 20^th^ bp downstream of the TTTR PAM (Fig. 4c). Consistently, Un1Cas12f1 exhibited three potential breaks at around 8^th^, 14^th^, and 25^th^ bp downstream of the TTTR PAM sequence (Fig. 4c). *As*Cas12f1 induced similar cleavage signals to Cas12a, approximately peaking at 14^th^ and 20^th^ bp, which was consistent with the previous report^10^ (Fig. 4c). Among these six tested Cas-nucleases, Tag-seq data revealed that *Sp*Cas9 was much less specific than Cas12a and Cas12f1, which triggered hundreds of off-target cleavages at other loci (Fig. 4d, e), indicating a weakness in targeting accuracy, and this conclusion was supported by the recently reported research that the Un1Cas121 and Cas12a had a higher editing safety^24^. Similar results were observed in MCF7, K562, and Jurkat cell lines (Fig. 4c-e and Supplementary Fig. 9-11).

**Figure 4.**
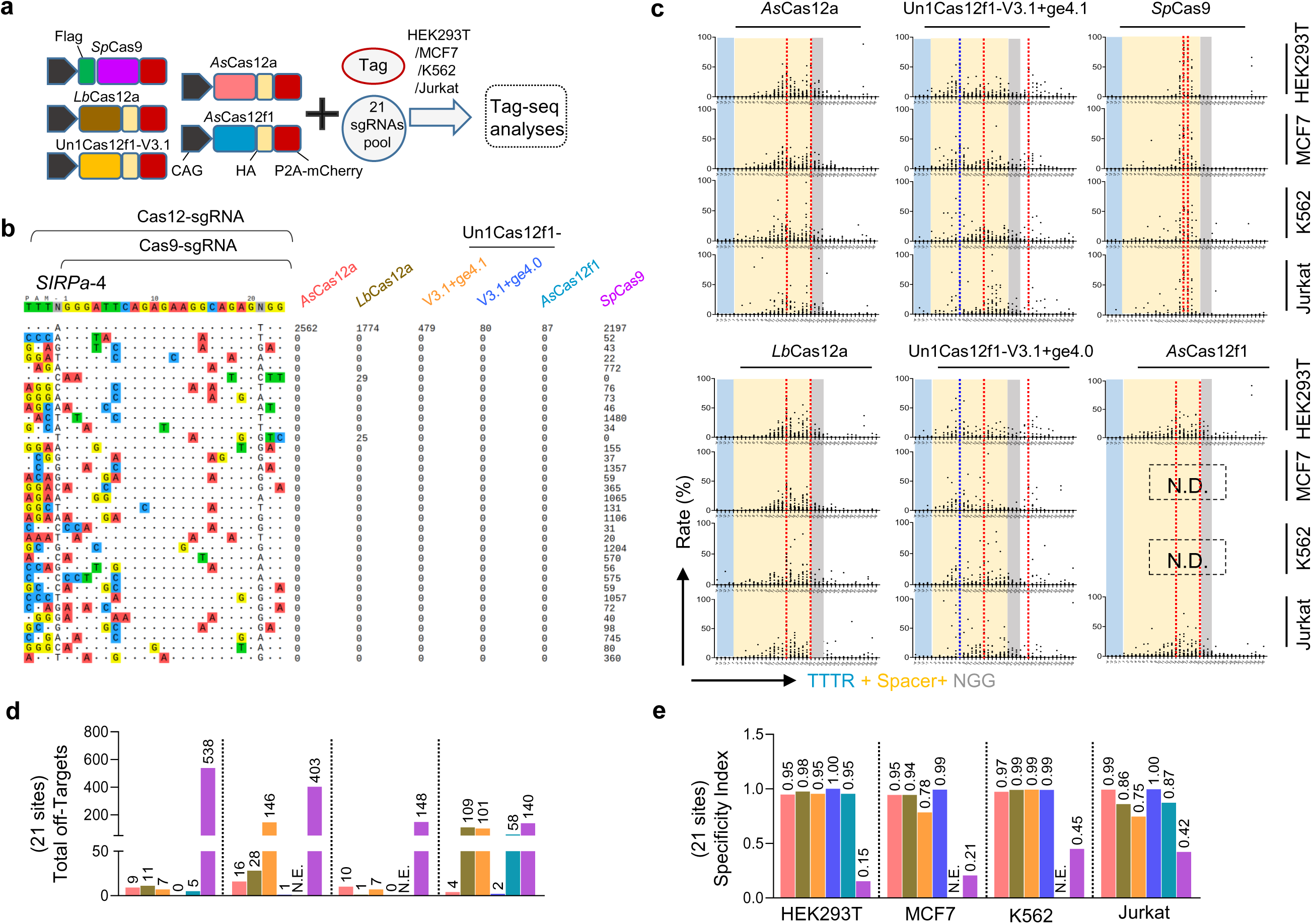
Editing specificities comparison of the DNA targeting CRISPR systems in human cells. **a** Schematic of the editing specificity analysis by Tag-seq among the DNA targeting CRISPR editors (CRISPR-*As*Cas12a, CRISPR-*Lb*Cas12a, CRISPR-Un1Cas12f1, CRISPR-*As*Cas12f1, and CRISPR-*Sp*Cas9 systems). **b** Tag-seq-based comparative analysis of DNA targeting CRISPR systems with twenty-one sgRNAs (also see Supplementary Fig. 9-12). The targeted sites for Cas12 and Cas9 share a common spacer sequence as shown at the top. As for Cas12a/Cas12f1, the sgRNA reference is the full sequence. As for SpCas9, the sgRNA sequence begins after the TTTR and ends with its NGG PAM. **c** The characteristics of the DNA targeting CRISPR systems induced DSBs revealed by Tag-seq. x-axis showing the location of the sgRNA. The red dotted line indicates the expected DSB sites, while the blue dotted line indicates the new potential breakpoint. **d** Total number of off-target sites detected with the twenty-one sgRNAs. N.E., no editing detected. **e** Specificity Index assessment (value was calculated by the ratio of total on-target reads to the on-target reads plus the off-target reads within the twenty-one sites).

The hypercompact size of Cas-nuclease provides great promise for programmable DNA modulation, especially for *in vivo* therapeutic applications. The Cas12f1 nuclease is one of these miniature enzymes, which can be easily packaged in an AAV vector for gene therapy delivery^9^. However, in terms of cleaving genomic DNA, our data and the recently reported study^24^ together revealed that this editor’s efficiency was generally lower than that of the widely used Cas9 and Cas12a (Fig. 3b, c). Therefore, to promote and extend the applications of the miniature Cas12f1 nucleases in genomic cleavage, further engineering aiming to increase the activity of Un1Cas12f1, as well as the *As*Cas12f1, is an urgent requirement. On the other hand, we also examined the binding ability by using DNase-inactive Un1Cas12f1 (dUn1Cas12f1) fusing transcription activator VPR to activate endogenous genes. Consistent with the original report^8^, our results showed that as an activator, the Un1Cas12f1 nuclease could induce endogenous gene robust expression with a comparable level to these of *Sp*Cas9 and Cas12a (Fig. 2), demonstrating its potential for gene-regulated therapy. Therefore, we highly recommend using these DNA-targeting editors according to the fitness of their unique properties to the intended scenarios. Generally, in a study that is needed to disrupt genes in cell lines or animal models, we will recommend the wild-type *Sp*Cas9 enzyme since the *Sp*Cas9 is still the most active nuclease to date and the activity is generally a priority over the specificity in these experiments. In contrast, for the experiments related to disrupting genes for therapy purposes, we will recommend the Cas12a effectors since the specificity (and thus the safety) is a priority over the activity and the Cas12a nucleases retain a relatively high targeting accuracy and also displays efficient editing ability. As for Cas12f1, we will recommend it to be used as an approach for gene activation relative therapy since it has a small size that can be easily packaged in the AAV vector and enable to induce robust gene activation. For more detailed comparisons among these DNA-targeting editors see Table 1.

**Table 1.** Comparison of the commonly used DNA targeting CRISPR editors

| **Editors**<br>**Contents** | **Cas9** | **Cas12a** | **Un1Cas12f1** | **AsCas12f1** |
| --- | --- | --- | --- | --- |
| PAM | NGG | TTTV | TTTR | NTTR |
| Size | SpCas9 (1368 aa) | AsCas12a (1307 aa);<br>LbCas12a (1228 aa) | 529 aa | 422 aa |
| Cutting site | -3<sup>th</sup>, -4<sup>th</sup> bp | 14<sup>th</sup>, 20<sup>th</sup> | 8<sup>th</sup>, 14<sup>th</sup>, and 25<sup>th</sup> | 14<sup>th</sup>, 20<sup>th</sup> |
| Break site | blunt end | sticky end | sticky end | sticky end |
| Specificity | * | *** | *** | *** |
| Activity | *** | ** | * | * |
| CRISPR-based<br>activation | *** | *** | *** | - |
| Application<br>scenarios | Disrupt genes in cells<br>or animal models | Disrupt genes for<br>therapy purposes | Gene-activation-based therapy | Need to improve |

Note: V=A/G/C; R=A/G; N=A/T/C/G.

## Conclusion

In summary, the parallel comparison of the commonly used DNA-targeting CRISPR editors provides basic information for the gene-editing toolkit, which can be a guideline and a helpful resource for this field.

## Acknowledgments

We are grateful to all members of the Lin labs for helpful comments and discussions on the manuscript.

## Methods

### Plasmids construction

All the Cas-nucleases expressing plasmids used in this study were constructed by standard PCR and molecular cloning into a plasmid containing a CAG promoter, a Flag/3xHA tag, a Cas-protein CDS expression cassette, and a P2A-mcherry reporter via Gibson Assembly. Except for the *Sp*Cas9 plasmid where the Flag tag was designed at its N-terminus, other vectors of Cas-protein used the 3xHA tags were constructed at their C-terminus. All sgRNAs targeting the genes of interest were designed through https://benchling.com/. sgRNA expression plasmids were constructed via digesting the sgRNA backbone plasmids that contained a human U6 promoter and a Cas-nuclease’s corresponding sgRNA scaffold sequences using BsmB I or Bbs I endonuclease (NEB) and then ligated the oligonucleotide duplexes into this cut backbone. The Cas-nuclease’s corresponding sgRNA scaffold sequences were listed in the Supplementary Information. All the plasmids were confirmed by Sanger sequencing and all the sgRNAs oligonucleotides used in this study were shown in Supplementary information Table S2.

### Cell culture

HEK293T and B16 cells were maintained in Dulbecco’s Modified Eagle’s Medium (DMEM, Life Technologies) at 37°C in a 5% CO2 humidified incubator. MCF7, Jurkat, and K562 cells were maintained in RPMI 1640 medium (Life Technologies) at 37°C in a 5% CO2 humidified incubator. All growth media were supplemented with 2 mM L-glutamine (Life Technologies), 100 U/mL penicillin, 100 µg/mL streptomycin (Life Technologies), and 10% fetal bovine serum. All the cell lines in this study were cultured in no more than 10 passages.

### Cell transfection and genomic DNA extraction

Generally, the HEK293T, MCF7, and B16 cells were transfected by the PEI method. For each transfection, approximately 2.0×10^5^/5.0×10^5^/1.0×10^6^cells were seeded in the 24/12/6-well plate, and the next day when cells grew up to 60%∼70%, the transfection was performed by PEI reagent with 250/500/1000 ng of Cas-nuclease expression plasmid and 250/500/1000 ng of the sgRNA-encoding plasmid per well in a 24/12/6-well plate. For the Jurkat and K562 cells, approximately 1.0×10^6^cells were electroporated by the Lonza Nucleofector Kit method with 1200 ng of Cas-nuclease expression plasmid and 800 ng of the sgRNA-encoding plasmids for each test. All cells in each well were harvested 3 days post-transfection and the genomic DNA was extracted by TIANamp Genomic DNA Kit (TIANGEN Biotech Co., Ltd., Beijing, China) following the manufacturer’s instructions.

### Quantitative real-time PCR

Total RNA from the transfected cells was isolated using Trizol Reagent (Thermo Fisher, USA) following the manufacturer’s instructions. Total RNA (1 μg) was reverse transcribed into cDNA and then quantitative real-time PCR was performed using a LightCycler 96 System (Roche, Switzerland). Relative gene expression was calculated using the 2^-ΔΔCt^ method after normalizing to GAPDH expression. The activation sgRNA used in this study and the qPCR primers were listed in Supplementary Table S2.

### Tag-seq analysis

Tag-seq can parallelly profile the off-target cleavages induced by Cas-nuclease at various sites in a single experiment and thus is a convenient and cost-efficient method for comparing the specificity among different nucleases, which was performed and described as previously reported^13, 27^. Briefly, for the HEK293T and MCF7, ∼6.0×10^5^ cells were transfected by PEI with 10 pmol Tag, 600 ng of Cas nuclease, and 600 ng pooled sgRNAs (21 guides) per well in a 12-well plate. For the Jurkat and K562, ∼1.0×10^6^ cells were transfected by Amaxa Cell Line Nucleofector Kit V (VCA-1003, Lonza, Switzerland) following the manufacturer’s instructions (2D) with 20 pmol Tag, 1200 ng of Cas nuclease, and 1000 ng pool sgRNAs (21 guides). Three days post-transfection genomic DNA was extracted for one-step libraries preparation by the Fragmentation, End Preparation, and dA-Tailing Module and Adapter Ligation Module kit (Vazyme Biotech Co., Ltd., Nanjing, China). The L and R libraries were constructed by PCR with library preparation primers, which were followed by sequencing (NovaSeq platform, Novogene, Beijing, China) and analysis with a Tag-seq bioinformatics pipeline. To profile the off-target effects induced by the Cas-nuclease at 21 sites in parallel, the 21 sgRNA sequences were parallelly listed in a format of “sgRNA-name TTTNN20NGG TTTN” in the “sgrna. lst” file and the parameter in “config. docker. text” file was set as follows: # parameter for detecting potential cutting sites MinSupportReadCount, 1; MinCuttingEventCount, 2 # off-target detection parameters MaxMismatch, 6; MaxGap, 1; MaxGapMismatch, 3.

The Tag-seq pipeline is available at https://github.com/zhoujj2013/Tag-seq and https://doi.org/10.5281/zenodo.4679460.

### Deep-seq analysis

Deep-seq was used to assess the editing activities of the DNA-targeting CRISPR editors. For HEK293T and MCF7, cells were transfected by PEI with 600 ng of Cas nuclease, and 600 ng pool sgRNAs (21 guides) per well in a 12-well plate. For the Jurkat and K562, cells were transfected by Amaxa Cell Line Nucleofector Kit V (VCA-1003, LONZA, Switzerland) following the manufacturer’s instructions (2D) with 1200 ng of Cas nuclease, and 1000 pool sgRNAs (21 guides). Two days post-transfection genomic DNA was extracted for deep-seq libraries preparation. Briefly, the primers with forward and reverse indexes were used to amplify the genomic regions in the first-round PCR. Then, equal amounts of the first PCR products were mixed and subjected to a second round of PCR with the P5- and P7-containing primers to generate the sequencing libraries. Paired-end sequencing was performed using the NovaSeq platform (Novogene, Beijing, China). Indel frequency was calculated as the ratio of (read counts with indel sequence)/(total sequencing read counts). The deep-seq primers were listed in Supplementary information Table S3.

### Western blotting

For detecting the expression of the Cas-proteins, HEK293T cells were transfected with the Cas-protein encoding plasmids using the PEI method. 2 days post-transfection cells were harvested and lysed in a 2×SDS loading buffer, and then boiled for 10 min. Lysates were resolved through SDS/PAGE and transferred onto a nitrocellulose membrane which was blocked using 5% non-fat milk and sequentially incubated with primary antibodies (anti-HA or anti-Flag, sigma, USA, anti-GADPH, Proteintech, China) and an HRP-conjugated horse anti-mouse IgG secondary antibody (CST, USA). All the probed proteins were finally detected through chemiluminescence following the manufacturer’s instructions (Pierce, USA).

### FACS analysis

All flow cytometry analyses were performed using FlowJo software (TreeStar, USA). To detect transfection efficiency, cells were harvested at the ending time and were determined as the proportion of the mCherry-positive (Cas-protein-P2A-mCherry cassette). To compare the editing activities of the DNA-targeting editors, cells were harvested two days post-transfection and the cleavage efficiency was determined as the proportion of GFP negative cells within the Cas-nucleases transfected cells (mCherry-positive).

### Activity and specificity Scoring

For the comparison of the performance among CRISPR-Cas systems, Tag-seq reads were used for calculating the targeting specificity and the editing activity, which was analyzed similarly to our previous study^27^. For the engineered CRISPR-Un1Cas12a systems comparison, editing activity scores were calculated as the mean ratio of the on-target reads across all the tested sites, normalized to the V3.1+ ge4.1 combination. The specificity Index was calculated as the ratio of the on-target reads to the on-target reads plus the off-target reads across all the tested sites.

### Data availability

All the next-generation sequencing data related to this study have been deposited in NCBI (Bioproject PRJNA799250). And other data that support the findings of this study are available from the corresponding author upon reasonable request.

## Author Contributions

Y.L. and H.H. conceived the study, designed the experiments, analyzed the data, and wrote the manuscript. H.H. performed most experiments. G.H. performed the Western bolting experiment. W.L., J.L., T.H., S.M., X.Z., and L.H. helped H.H. with some experiments. Z.T. and Y.H. helped to perform bioinformatics analyses.

## Supplementary Information

### DNA sequences

#### l Expression of the ge3.0 sgRNA

Red: U6 promoter, blue: ge3 scaffold

gagggcctatttcccatgattccttcatatttgcatatacgatacaaggctgttagagagataattagaattaatttgactgtaaacacaaagatattagtacaaaat acgtgacgtagaaagtaataatttcttgggtagtttgcagttttaaaattatgttttaaaatggactatcatatgcttaccgtaacttgaaagtatttcgatttcttggcttt atatatcttgtggaaaggacgaaacaccgACCGCTTCACCAAAAGCTGTCCCTTAGGGGATTAGAACTTGAGTGAAGGTG GGCTGCTTGCATCAGCCTAATGTCGAGAAGTGCTTTCTTCGGAAAGTAACCCTCGAAACAAATTCAGTGCTC CTCTCCAATTCTGCACAAGAAAGTTGCAGAACCCGAATAGAGCAATGAAGGAATGCAAC/20bp-oligo/ttttatttttt

#### l Expression of the ge4.0 sgRNA

Red: U6 promoter, blue: ge4 scaffold

gagggcctatttcccatgattccttcatatttgcatatacgatacaaggctgttagagagataattagaattaatttgactgtaaacacaaagatattagtacaaaat acgtgacgtagaaagtaataatttcttgggtagtttgcagttttaaaattatgttttaaaatggactatcatatgcttaccgtaacttgaaagtatttcgatttcttggcttt atatatcttgtggaaaggacgaaacaccgACCGCTTCACCAAAAGCTGTCCCTTAGGGGATTAGAACTTGAGTGAAGGTG GGCTGCTTGCATCAGCCTAATGTCGAGAAGTGCTTTCTTCGGAAAGTAACCCTCGAAACAAAgaaaGGAATGC AAC/20bp-oligo/ttttatttttt

#### l Expression of the ge4.1 sgRNA

Red: U6 promoter, blue: ge4.1 scaffold

gagggcctatttcccatgattccttcatatttgcatatacgatacaaggctgttagagagataattagaattaatttgactgtaaacacaaagatattagtacaaaat acgtgacgtagaaagtaataatttcttgggtagtttgcagttttaaaattatgttttaaaatggactatcatatgcttaccgtaacttgaaagtatttcgatttcttggcttt atatatcttgtggaaaggacgaaacaccgACCGCTTCACTTAGAGTGAAGGTGGGCTGCTTGCATCAGCCTAATGTCGAG AAGTGCTTTCTTCGGAAAGTAACCCTCGAAACAAAgaaaGGAATGCAAC/20bp-oligo/ttttatttttt

#### • Expression of the AsCas12f1 sgRNA

Red: U6 promoter, blue: AsCas12f1 sgRNA scaffold gagggcctatttcccatgattccttcatatttgcatatacgatacaaggctgttagagagataattagaattaatttgactgtaaacacaaagatattagtacaaaat acgtgacgtagaaagtaataatttcttgggtagtttgcagttttaaaattatgttttaaaatggactatcatatgcttaccgtaacttgaaagtatttcgatttcttggcttt atatatcttgtggaaaggacgaaacaccgATTCGTCGGTTCAGCGACGATAAGCCGAGAAGTGCCAATAAAACTGTTAAGT GGTTTGGTAACGCTCGGTAAGGTAGCCAAAAGGCTGAAACTCCGTGCACAAAGACCGCACGGACGCTTCAC ATATAGCTCATAAACAAGGGTTTGCGAGCTAGCTTGTGGAGTGTGAAC/20bp-oligo/tttttt

#### • Expression of the LbCas12a sgRNA

Red: U6 promoter, blue: LbCas12a sgRNA scaffold gagggcctatttcccatgattccttcatatttgcatatacgatacaaggctgttagagagataattagaattaatttgactgtaaacacaaagatattagtacaaaat acgtgacgtagaaagtaataatttcttgggtagtttgcagttttaaaattatgttttaaaatggactatcatatgcttaccgtaacttgaaagtatttcgatttcttggcttt atatatcttgtggaaaggacgaaacaccgAATTTCTACTAAGTGTAGAT/20bp-oligo/ttttatttttt

#### • Expression of the AsCas12a sgRNA

Red: U6 promoter, blue: AsCas12a sgRNA scaffold gagggcctatttcccatgattccttcatatttgcatatacgatacaaggctgttagagagataattagaattaatttgactgtaaacacaaagatattagtacaaaat acgtgacgtagaaagtaataatttcttgggtagtttgcagttttaaaattatgttttaaaatggactatcatatgcttaccgtaacttgaaagtatttcgatttcttggcttt atatatcttgtggaaaggacgaaacaccgAATTTCTACTCTTGTAGAT/20bp-oligo/ttttatttttt

#### l Expression of the Cas9 FE2.1 sgRNA

Red: U6 promoter, blue: Cas9 FE2.1 scaffold

gagggcctatttcccatgattccttcatatttgcatatacgatacaaggctgttagagagataattagaattaatttgactgtaaacacaaagatattagtacaaaat acgtgacgtagaaagtaataatttcttgggtagtttgcagttttaaaattatgttttaaaatggactatcatatgcttaccgtaacttgaaagtatttcgatttcttggcttt atatatcttgtggaaaggacgaaacacc/g+19bp-oligo/GTTTAAGAGCTATGCTGGAAACAGCATAGCAAGTTTAAATAAGG CTAGTCCGTTATCAACTTTGCTGGAAACAGCAAAGTGGCACCGAGTCGGTGCTTTTTTT

#### • Expression of the AsCas12f1 nuclease

Orange: SV40 NLS, blue: AsCas12f1, violet: nucleoplasmin NLS, green: 3XHA, yellow: P2A, Red: mCherry CCAAAGAAGAAGCGGAAGGTCATGATCAAGGTGTACAGATACGAGATCGTGAAGCCTCTGGACCTGGACTG GAAGGAGTTCGGCACCATCCTGAGACAGCTGCAGCAGGAAACCAGATTCGCCCTGAATAAGGCCACACAGC TGGCCTGGGAGTGGATGGGCTTCAGCAGCGACTACAAGGATAACCACGGCGAGTACCCCAAGAGCAAGGA CATCCTGGGCTACACCAACGTGCACGGCTACGCCTACCACACCATCAAGACAAAGGCCTACAGACTGAACTCTGGAAATCTGAGCCAGACCATCAAGAGAGCCACAGACAGGTTCAAGGCCTACCAGAAGGAGATCCTGCGCGGCGACATGTCTATCCCCAGCTACAAGAGGGACATCCCCCTGGACCTGATCAAGGAGAACATCTCCGTGAACAGGATGAATCACGGCGACTACATCGCCAGCCTGTCTCTGCTGAGCAACCCCGCCAAGCAGGAGATGAACGTGAAGAGAAAGATCTCCGTGATCATCATCGTGAGGGGCGCCGGCAAGACCATCATGGACAGAATCCTGTCCGGCGAGTACCAGGTGAGCGCCAGCCAGATTATCCACGACGACCGGAAGAACAAGTGGTACCTGAACATCAGCTACGACTTCGAGCCACAGACCAGAGTGCTGGACCTGAACAAGATCATGGGCATTGACCTGGGCGTGGCCGTGGCCGTGTACATGGCCTTCCAGCACACCCCCGCCAGGTACAAGCTGGAGGGCGGCGAGATTGAGAACTTCAGGAGGCAGGTGGAGAGCCGGCGCATCTCCATGCTGAGACAGGGCAAGTACGCCGGCGGCGCCAGGGGCGGCCACGGCAGAGACAAGAGAATCAAGCCCATTGAGCAGCTGAGGGATAAGATCGCCAATTTCAGAGACACCACCAATCACCGGTACAGCAGATACATCGTGGACATGGCCATCAAGGAGGGCTGCGGCACAATCCAGATGGAGGATCTGACAAACATCAGAGACATCGGCAGCAGATTCCTGCAGAACTGGACCTACTACGACCTGCAGCAGAAGATCATCTACAAGGCCGAGGAGGCCGGCATCAAAGTGATCAAGATCGACCCCCAGTACACCAGCCAGAGATGCTCCGAGTGCGGCAACATCGACTCCGGCAACAGAATCGGCCAGGCCATCTTTAAGTGCCGGGCCTGCGGCTACGAGGCCAACGCCGACTACAACGCCGCCCGGAATATCGCCATCCCCAACATCGACAAGATCATCGCCGAGAGCATTAAGAAAAGGCCGGCGGCCACGAAAAAGGCCGGCCAGGCAAAAAAGAAAAAGGGATCCTACCCATACGATGTTCCAGATTACGCTTATCCCTACGACGTGCCTGATTATGCATACCCATACGATGTCCCCGACTATGCCCTCGAGAGCACCGGTGGCAGCGGAGCTACTAACTTCAGCCTGCTGAAGCAGGCTGGAGACGTGGAGGAGAACCCTGGACCTGCCGGTATGGTGAGCAAGGGCGAGGAGGATAACATGGCCATCATCAAGGAGTTCATGCGCTTCAAGGTGCACATGGAGGGCTCCGTGAACGGCCACGAGTTCGAGATCGAGGGCGAGGGCGAGGGCCGCCCCTACGAGGGCACCCAGACCGCCAAGCTGAAGGTGACCAAGGGTGGCCCCCTGCCCTTCGCCTGGGACATCCTGTCCCCTCAGTTCATGTACGGCTCCAAGGCCTACGTGAAGCACCCCGCCGACATCCCCGACTACTTGAAGCTGTCCTTCCCCGAGGGCTTCAAGTGGGAGCGCGTGATGAACTTCGAGGACGGCGGCGTGGTGACCGTGACCCAGGACTCCTCCCTGCAGGACGGCGAGTTCATCTACAAGGTGAAGCTGCGCGGCACCAACTTCCCCTCCGACGGCCCCGTAATGCAGAAGAAGACCATGGGCTGGGAGGCCTCCTCCGAGCGGATGTA CCCCGAGGACGGCGCCCTGAAGGGCGAGATCAAGCAGAGGCTGAAGCTGAAGGACGGCGGCCACTACGA CGCTGAGGTCAAGACCACCTACAAGGCCAAGAAGCCCGTGCAGCTGCCCGGCGCCTACAACGTCAACATC AAGTTGGACATCACCTCCCACAACGAGGACTACACCATCGTGGAACAGTACGAACGCGCCGAGGGCCGCC ACTCCACCGGCGGCATGGACGAGCTGTACAAGTAA

#### • Expression of the Un1Cas12f1-WT nuclease

Orange: SV40 NLS, blue: Un1Cas12f1-WT, violet: nucleoplasmin NLS, green: 3XHA, yellow: P2A, Red: mCherry CCAAAGAAGAAGCGGAAGGTCGGTATCCACGGAGTCCCAGCAGCCATGGCCAAGAACACAATTACAAAGAC ACTGAAGCTGAGGATCGTGAGACCATACAACAGCGCTGAGGTCGAGAAGATTGTGGCTGATGAAAAGAACA ACAGGGAAAAGATCGCCCTCGAGAAGAACAAGGATAAGGTGAAGGAGGCCTGCTCTAAGCACCTGAAAGTGGCCGCCTACTGCACCACACAGGTGGAGAGGAACGCCTGTCTGTTTTGTAAAGCTCGGAAGCTGGATGATAAGTTTTACCAGAAGCTGCGGGGCCAGTTCCCCGATGCCGTCTTTTGGCAGGAGATTAGCGAGATCTTCAGACAGCTGCAGAAGCAGGCCGCCGAGATCTACAACCAGAGCCTGATCGAGCTCTACTACGAGATCTTCATCAAGGGCAAGGGCATTGCCAACGCCTCCTCCGTGGAGCACTACCTGAGCGACGTGTGCTACACAAGAGCCGCCGAGCTCTTTAAGAACGCCGCTATCGCTTCCGGGCTGAGGAGCAAGATTAAGAGTAACTTCCGGCTCAAGGAGCTGAAGAACATGAAGAGCGGCCTGCCCACTACAAAGAGCGACAACTTCCCAATTCCACTGGTGAAGCAGAAGGGGGGCCAGTACACAGGGTTCGAGATTTCCAACCACAACAGCGACTTTATTATTAAGATCCCCTTTGGCAGGTGGCAGGTCAAGAAGGAGATTGACAAGTACAGGCCCTGGGAGAAGTTTGATTTCGAGCAGGTGCAGAAGAGCCCCAAGCCTATTTCCCTGCTGCTGTCCACACAGCGGCGGAAGAGGAACAAGGGGTGGTCTAAGGATGAGGGGACCGAGGCCGAGATTAAGAAAGTGATGAACGGCGACTACCAGACAAGCTACATCGAGGTCAAGCGGGGCAGTAAGATTTGCGAGAAGAGCGCCTGGATGCTGAACCTGAGCATTGACGTGCCAAAGATTGATAAGGGCGTGGACCCCAGCATCATCGGAGGGATCGATGTGGGGGTCAAGAGCCCCCTCGTGTGCGCCATCAACAACGCCTTCAGCAGGTACAGCATCTCCGATAACGACCTGTTCCACTTTAACAAGAAGATGTTCGCCCGGCGGAGGATTTTGCTCAAGAAGAACCGGCACAAGCGGGCCGGACACGGGGCCAAGAACAAGCTCAAGCCCATCACTATCCTGACCGAGAAGAGCGAGAGGTTCAGGAAGAAGCTCATCGAGAGATGGGCCTGCGAGATCGCCGATTTCTTTATTAAGAACAAGGTCGGAACAGTGCAGATGGAGAACCTCGAGAGCATGAAGAGGAAGGAGGATTCCTACTTCAACATTCGGCTGAGGGGGTTCTGGCCCTACGCTGAGATGCAGAACAAGATTGAGTTTAAGCTGAAGCAGTACGGGATTGAGATCCGGAAGGTGGCCCCCAACAACACCAGCAAGACCTGCAGCAAGTGCGGGCACCTCAACAACTACTTCAACTTCGAGTACCGGAAGAAGAACAAGTTCCCACACTTCAAGTGCGAGAAGTGCAACTTTAAGGAGAACGCCGATTACAACGCCGCCCTGAACATCAGCAACCCTAAGCTGAAGAGCACTAAGGAGGAGCCCAAAAGGCCGGCGGCCACGAAAAAGGCCGGCCAGGCAAAAAAGAAAAAGGGATCCTACCCATACGATGTTCCAGATTACGCTTATCCCTACGACGTGCCTGATTATGCATACCCATACGATGTCCCCGACTATGCCCTCGAGAGCACCGGTGGCAGCGGAGCTACTAACTTCAGCCTGCTGAAGCAGGCTGGAGACGTGGAGGAGAACCCTGGACCTGCCGGTATGGTGAGCAAGGGCGAGGAGGATAACATGGCCATCATCAAGGAGTTCATGCGCTTCAAGGTGCACATGGAGGGCTCCGTGAACGGCCACGAGTTCGAGATCGAGGGCGAGGGCGAGGGCCGCCCCTACGAGGGCACCCAGACCGCCAAGCTGAAGGTGACCAAGGGTGGCCCCCTGCCCTTCGCCTGGGACATCCTGTCCCCTCAGTTCATGTACGGCTCCAAGGCCTACGTGAAGCACCCCGCCGACATCCCCGACTACTTGAAGCTGTCCTTCCCCGAGGGCTTCAAGTGGGAGCGCGTGATGAACTTCGAGGACGGCGGCGTGGTGACCGTGACCCAGGACTCCTCCCTGCAGGACGGCGAGTTCATCTACAAGGTGAAGCTGCGCGGCACCAACTTCCCCTCCGACGGCCCCGTAATGCAGAAGAAGACCATGGGCTGGGAGGCCTCCTCCGAGCGGATGTACCCCGAGGACGGCGCCCTGAAGGGCGAGATCAAGCAGAGGCTGAAGCTGAAGGACGGCGGCCACTACGACGCTGAGGTCAAGACCACCTACAAGGCCAAGAAGCCCGTGCAGCTGCCCGGCGCCTACAACGTCAACATCAAGTTGGACATC ACCTCCCACAACGAGGACTACACCATCGTGGAACAGTACGAACGCGCCGAGGGCCGCCACTCCACCGGCG GCATGGACGAGCTGTACAAGTAA

#### • Expression of the Un1Cas12f1-V3.1 nuclease

Orange: SV40 NLS, blue: Un1Cas12f1-V3.1(D143R/T147R/E151A), violet: nucleoplasmin NLS, green: 3XHA, yellow: P2A, Red: mCherry CCAAAGAAGAAGCGGAAGGTCGGTATCCACGGAGTCCCAGCAGCCATGGCCAAGAACACAATTACAAAGAC ACTGAAGCTGAGGATCGTGAGACCATACAACAGCGCTGAGGTCGAGAAGATTGTGGCTGATGAAAAGAACA ACAGGGAAAAGATCGCCCTCGAGAAGAACAAGGATAAGGTGAAGGAGGCCTGCTCTAAGCACCTGAAAGTG GCCGCCTACTGCACCACACAGGTGGAGAGGAACGCCTGTCTGTTTTGTAAAGCTCGGAAGCTGGATGATAA GTTTTACCAGAAGCTGCGGGGCCAGTTCCCCGATGCCGTCTTTTGGCAGGAGATTAGCGAGATCTTCAGAC AGCTGCAGAAGCAGGCCGCCGAGATCTACAACCAGAGCCTGATCGAGCTCTACTACGAGATCTTCATCAAG GGCAAGGGCATTGCCAACGCCTCCTCCGTGGAGCACTACCTGAGC<u>AGA</u>GTGTGCTAC<u>AGA</u>AGAGCCGCC<u>G</u> <u>CT</u>CTCTTTAAGAACGCCGCTATCGCTTCCGGGCTGAGGAGCAAGATTAAGAGTAACTTCCGGCTCAAGGAGC TGAAGAACATGAAGAGCGGCCTGCCCACTACAAAGAGCGACAACTTCCCAATTCCACTGGTGAAGCAGAAGGGGGGCCAGTACACAGGGTTCGAGATTTCCAACCACAACAGCGACTTTATTATTAAGATCCCCTTTGGCAGGTGGCAGGTCAAGAAGGAGATTGACAAGTACAGGCCCTGGGAGAAGTTTGATTTCGAGCAGGTGCAGAAGAGCCCCAAGCCTATTTCCCTGCTGCTGTCCACACAGCGGCGGAAGAGGAACAAGGGGTGGTCTAAGGATGAGGGGACCGAGGCCGAGATTAAGAAAGTGATGAACGGCGACTACCAGACAAGCTACATCGAGGTCAAGCGGGGCAGTAAGATTTGCGAGAAGAGCGCCTGGATGCTGAACCTGAGCATTGACGTGCCAAAGATTGATAAGGGCGTGGACCCCAGCATCATCGGAGGGATCGATGTGGGGGTCAAGAGCCCCCTCGTGTGCGCCATCAACAACGCCTTCAGCAGGTACAGCATCTCCGATAACGACCTGTTCCACTTTAACAAGAAGATGTTCGCCCGGCGGAGGATTTTGCTCAAGAAGAACCGGCACAAGCGGGCCGGACACGGGGCCAAGAACAAGCTCAAGCCCATCACTATCCTGACCGAGAAGAGCGAGAGGTTCAGGAAGAAGCTCATCGAGAGATGGGCCTGCGAGATCGCCGATTTCTTTATTAAGAACAAGGTCGGAACAGTGCAGATGGAGAACCTCGAGAGCATGAAGAGGAAGGAGGATTCCTACTTCAACATTCGGCTGAGGGGGTTCTGGCCCTACGCTGAGATGCAGAACAAGATTGAGTTTAAGCTGAAGCAGTACGGGATTGAGATCCGGAAGGTGGCCCCCAACAACACCAGCAAGACCTGCAGCAAGTGCGGGCACCTCAACAACTACTTCAACTTCGAGTACCGGAAGAAGAACAAGTTCCCACACTTCAAGTGCGAGAAGTGCAACTTTAAGGAGAACGCCGATTACAACGCCGCCCTGAACATCAGCAACCCTAAGCTGAAGAGCACTAAGGAGGAGCCCAAAAGGCCGGCGGCCACGAAAAAGGCCGGCCAGGCAAAAAAGAAAAAGGGATCCTACCCATACGATGTTCCAGATTACGCTTATCCCTACGACGTGCCTGATTATGCATACCCATACGATGTCCCCGACTATGCCCTCGAGAGCACCGGTGGCAGCGGAGCTACTAACTTCAGCCTGCTGAAGCAGGCTGGAGACGTGGAGGAGAACCCTGGACCTGCCGGTATGGTGAGCAAGGGCGAGGAGGATAACATGGCCATCATCAAGGAGTTCATGCGCTTCAAGGTGCACATGGAGGGCTCCGTGAACGGCCACGAGTTCGAGATCGAGGGCGAGGGCGAGGGCCGCCCCTACGAGGGCACCCAGACCGCCAAGCTGAAGGTGACCAAGGGTGGCCCCCTGCCCTTCGCCTGGGACATCCTGTCCCCTCAGTTCATGTACGGCTCCAAGGCCTACGTGAAGCACCCCGCCGACATCCCCGACTACTTGAAGCTGTCCTTCCCCGAGGGCTTCAAGTGGGAGCGCGTGATGAACTTCGAGGACGGCGGCGTGGTGACCGTGACCCAGGACTCCTCCCTGCAGGACGGCGAGTTCATCTACAAGGTGAAGCTGCGCGGCACCAACTTCCCCTCCGACGGCCCCGTAATGCAGAAGAAGACCATGGGCTGGGAGGCCTCCTCCGAGCGGATGTACCCCGAGGACGGCGCC CTGAAGGGCGAGATCAAGCAGAGGCTGAAGCTGAAGGACGGCGGCCACTACGACGCTGAGGTCAAGACC ACCTACAAGGCCAAGAAGCCCGTGCAGCTGCCCGGCGCCTACAACGTCAACATCAAGTTGGACATCACCTC CCACAACGAGGACTACACCATCGTGGAACAGTACGAACGCGCCGAGGGCCGCCACTCCACCGGCGGCATG GACGAGCTGTACAAGTAA

#### • Expression of the AsCas12a nuclease

Blue: AsCas12a, violet: nucleoplasmin NLS, green: 3XHA, yellow: P2A, Red: mCherry ATGACACAGTTCGAGGGCTTTACCAACCTGTATCAGGTGAGCAAGACACTGCGGTTTGAGCTGATCCCACA GGGCAAGACCCTGAAGCACATCCAGGAGCAGGGCTTCATCGAGGAGGACAAGGCCCGCAATGATCACTACAAGGAGCTGAAGCCCATCATCGATCGGATCTACAAGACCTATGCCGACCAGTGCCTGCAGCTGGTGCAGCTGGATTGGGAGAACCTGAGCGCCGCCATCGACTCCTATAGAAAGGAGAAAACCGAGGAGACAAGGAACGCCCTGATCGAGGAGCAGGCCACATATCGCAATGCCATCCACGACTACTTCATCGGCCGGACAGACAACCTGACCGATGCCATCAATAAGAGACACGCCGAGATCTACAAGGGCCTGTTCAAGGCCGAGCTGTTTAATGGCAAGGTGCTGAAGCAGCTGGGCACCGTGACCACAACCGAGCACGAGAACGCCCTGCTGCGGAGCTTCGACAAGTTTACAACCTACTTCTCCGGCTTTTATGAGAACAGGAAGAACGTGTTCAGCGCCGAGGATATCAGCACAGCCATCCCACACCGCATCGTGCAGGACAACTTCCCCAAGTTTAAGGAGAATTGTCACATCTTCACACGCCTGATCACCGCCGTGCCCAGCCTGCGGGAGCACTTTGAGAACGTGAAGAAGGCCATCGGCATCTTCGTGAGCACCTCCATCGAGGAGGTGTTTTCCTTCCCTTTTTATAACCAGCTGCTGACACAGACCCAGATCGACCTGTATAACCAGCT GCTGGGAGGAATCTCTCGGGAGGCAGGCACCGAGAAGATCAAGGGCCTGAACGAGGTGCTGAATCTGGCCATCCAGAAGAATGATGAGACAGCCCACATCATCGCCTCCCTGCCACACAGATTCATCCCCCTGTTTAAGCAGATCCTGTCCGATAGGAACACCCTGTCTTTCATCCTGGAGGAGTTTAAGAGCGACGAGGAAGTGATCCAGTCCTTCTGCAAGTACAAGACACTGCTGAGAAACGAGAACGTGCTGGAGACAGCCGAGGCCCTGTTTAACGAGCTGAACAGCATCGACCTGACACACATCTTCATCAGCCACAAGAAGCTGGAGACAATCAGCAGCGCCCTGTGCGACCACTGGGATACACTGAGGAATGCCCTGTATGAGCGGAGAATCTCCGAGCTGACAGGCAAGATCACCAAGTCTGCCAAGGAGAAGGTGCAGCGCAGCCTGAAGCACGAGGATATCAACCTGCAGGAGATCATCTCTGCCGCAGGCAAGGAGCTGAGCGAGGCCTTCAAGCAGAAAACCAGCGAGATCCTGTCCCACGCACACGCCGCCCTGGATCAGCCACTGCCTACAACCCTGAAGAAGCAGGAGGAGAAGGAGATCCTGAAGTCTCAGCTGGACAGCCTGCTGGGCCTGTACCACCTGCTGGACTGGTTTGCCGTGGATGAGTCCAACGAGGTGGACCCCGAGTTCTCTGCCCGGCTGACCGGCATCAAGCTGGAGATGGAGCCTTCTCTGAGCTTCTACAACAAGGCCAGAAATTATGCCACCAAGAAGCCCTACTCCGTGGAGAAGTTCAAGCTGAACTTTCAGATGCCTACACTGGCCTCTGGCTGGGACGTGAATAAGGAGAAGAACAATGGCGCCATCCTGTTTGTGAAGAACGGCCTGTACTATCTGGGCATCATGCCAAAGCAGAAGGGCAGGTATAAGGCCCTGAGCTTCGAGCCCACAGAGAAAACCAGCGAGGGCTTTGATAAGATGTACTATGACTACTTCCCTGATGCCGCCAAGATGATCCCAAAGTGCAGCACCCAGCTGAAGGCCGTGACAGCCCACTTTCAGACCCACACAACCCCCATCCTGCTGTCCAACAATTTCATCGAGCCTCTGGAGATCACAAAGGAGATCTACGACCTGAACAATCCTGAGAAGGAGCCAAAGAAGTTTCAGACAGCCTACGCCAAGAAAACCGGCGACCAGAAGGGCTACAGAGAGGCCCTGTGCAAGTGGATCGACTTCACAAGGGATTTTCTGTCCAAGTATACCAAGACAACCTCTATCGATCTGTCTAGCCTGCGGCCATCCTCTCAGTATAAGGACCTGGGCGAGTACTATGCCGAGCTGAATCCCCTGCTGTACCACATCAGCTTCCAGAGAATCGCCGAGAAGGAGATCATGGATGCCGTGGAGACAGGCAAGCTGTACCTGTTCCAGATCTATAACAAGGACTTTGCCAAGGGCCACCACGGCAAGCCTAATCTGCACACACTGTATTGGACCGGCCTGTTTTCTCCAGAGAACCTGGCCAAGACAAGCATCAAGCTGAATGGCCAGGCCGAGCTGTTCTACCGCCCTAAGTCCAGGATGAAGAGGATGGCACACCGGCTGGGAGAGAAGATGCTGAACAAGAAGCTGAAGGATCAGAAAACCCCAATCCCCGACACCCTGTACCAGGAGCTGTACGACTATGTGAATCACAGACTGTCCCACGACCTGTCTGATGAGGCCAGGGCCCTGCTGCCCAACGTGATCACCAAGGAGGTGTCTCACGAGATCATCAAGGATAGGCGCTTTACCAGCGACAAGTTCTTTTTCCACGTGCCTATCACACTGAACTATCAGGCCGCCAATTCCCCATCTAAGTTCAACCAGAGGGTGAATGCCTACCTGAAGGAGCACCCCGAGACACCTATCATCGGCATCGATCGGGGCGAGAGAAACCTGATCTATATCACAGTGATCGACTCCACCGGCAAGATCCTGGAGCAGCGGAGCCTGAACACCATCCAGCAGTTTGATTACCAGAAGAAGCTGGACAACAGGGAGAAGGAGAGGGTGGCAGCAAGGCAGGCCTGGTCTGTGGTGGGCACAATCAAGGATCTGAAGCAGGGCTATCTGAGCCAGGTCATCCACGAGATCGTGGACCTGATGATCCACTACCAGGCCGTGGTGGTGCTGGAGAACCTGAATTTCGGCTTTAAGAGCAAGAGGACCGGCATCGCCGAGAAGGCCGTGTACCAGCAGTTCGAGAAGATGCTGATCGATAAGCTGAATTGCCTGGTGCTGAAGGACTATCCAGCAGAGAAAGTGGGAGGCGTGCTGAACCCATACCAGCTGACAGACCAGTTCACCTCCTTTGCCAAGATGGGCACCCAGTCTGGCTTCCTGTTTTACGTGCCTGCCCCATATACATCTAAGATCGATCCCCTGACCGGCTTCGTGGACCCCTTCGTGTGGAAAACCATCAAGAATCACGAGAGCCGCAAGCACTTCCTGGAGGGCTTCGACTTTCTGCACTACGACGTGAAAACCGGCGACTTCATCCTGCACTTTAAGATGAACAGAAATCTGTCCTTCCAGAGGGGCCTGCCCGGCTTTATGCCTGCATGGGATATCGTGTTCGAGAAGAACGAGACACAGTTTGACGCCAAGGGCACCCCTTTCATCGCCGGCAAGAGAATCGTGCCAGTGATCGAGAATCACAGATTCACCGGCAGATACCGGGACCTGTATCCTGCCAACGAGCTGATCGCCCTGCTGGAGGAGAAGGGCATCGTGTTCAGGGATGGCTCCAACATCCTGCCAAAGCTGCTGGAGAATGACGATTCTCACGCCATCGACACCATGGTGGCCCTGATCCGCAGCGTGCTGCAGATGCGGAACTCCAATGCCGCCACAGGCGAGGACTATATCAACAGCCCCGTGCGCGATCTGAATGGCGTGTGCTTCGACTCCCGGTTTCAGAACCCAGAGTGGCCCATGGACGCCGATGCCAATGGCGCCTACCACATCGCCCTGAAGGGCCAGCTGCTGCTGAATCACCTGAAGGAGAGCAAGGATCTGAAGCTGCAGAACGGCATCTCCAATCAGGACTGGCTGGCCTACATCCAGGAGCTGCGCAACAAAAGGCCGGCGGCCACGAAAAAGGCCGGCCAGGCAAAAAAGAAAAAGGGATCCTACCCATACGATGTTCCAGATTACGCTTATCCCTACGACGTGCCTGATTATGCATACCCATACGATGTCCCCGACTATGCCCTCGAGAGCACCGGTGGCAGCGGAGCTACTAACTTCAGCCTGCTGAAGCAGGCTGGAGACGTGGAGGAGAACCCTGGACCTGCCGGTATGGTGAGCAAGGGCGAGGAGGATAACATGGCCATCATCAAGGAGTTCATGCGCTTCAAGGTGCACATGGAGGGCTCCGTGAACGGCCACGAGTTCGAGATCGAGGGCGAGGGCGAGGGCC GCCCCTACGAGGGCACCCAGACCGCCAAGCTGAAGGTGACCAAGGGTGGCCCCCTGCCCTTCGCCTGGGACATCCTGTCCCCTCAGTTCATGTACGGCTCCAAGGCCTACGTGAAGCACCCCGCCGACATCCCCGACTACTTGAAGCTGTCCTTCCCCGAGGGCTTCAAGTGGGAGCGCGTGATGAACTTCGAGGACGGCGGCGTGGTGACCGTGACCCAGGACTCCTCCCTGCAGGACGGCGAGTTCATCTACAAGGTGAAGCTGCGCGGCACCAACTTCCCCTCCGACGGCCCCGTAATGCAGAAGAAGACCATGGGCTGGGAGGCCTCCTCCGAGCGGATGTACCCCGAGGACGGCGCCCTGAAGGGCGAGATCAAGCAGAGGCTGAAGCTGAAGGACGGCGGCCACTACGACGCTGAGGTCAAGACCACCTACAAGGCCAAGAAGCCCGTGCAGCTGCCCGGCGCCTACAACGTCAACATCAAGTT GGACATCACCTCCCACAACGAGGACTACACCATCGTGGAACAGTACGAACGCGCCGAGGGCCGCCACTCC ACCGGCGGCATGGACGAGCTGTACAAGTAA

#### • Expression of the LbCas12a nuclease

Blue: LbCas12a, violet: nucleoplasmin NLS, green: 3XHA, yellow: P2A, Red: mCherry ATGAGCAAGCTGGAGAAGTTTACAAACTGCTACTCCCTGTCTAAGACCCTGAGGTTCAAGGCCATCCCTGTG GGCAAGACCCAGGAGAACATCGACAATAAGCGGCTGCTGGTGGAGGACGAGAAGAGAGCCGAGGATTATAAGGGCGTGAAGAAGCTGCTGGATCGCTACTATCTGTCTTTTATCAACGACGTGCTGCACAGCATCAAGCTGAAGAATCTGAACAATTACATCAGCCTGTTCCGGAAGAAAACCAGAACCGAGAAGGAGAATAAGGAGCTGGAGAACCTGGAGATCAATCTGCGGAAGGAGATCGCCAAGGCCTTCAAGGGCAACGAGGGCTACAAGTCCCTGTTTAAGAAGGATATCATCGAGACAATCCTGCCAGAGTTCCTGGACGATAAGGACGAGATCGCCCTGGTGAACAGCTTCAATGGCTTTACCACAGCCTTCACCGGCTTCTTTGATAACAGAGAGAATATGTTTTCCGAGGAGGCCAAGAGCACATCCATCGCCTTCAGGTGTATCAACGAGAATCTGACCCGCTACATCTCTAATATGGACATCTTCGAGAAGGTGGACGCCATCTTTGATAAGCACGAGGTGCAGGAGATCAAGGAGAAGATCCTGAACAGCGACTATGATGTGGAGGATTTCTTTGAGGGCGAGTTCTTTAACTTTGTGCTGACACAGGAGGGCATCGACGTGTATAACGCCATCATCGGCGGCTTCGTGACCGAGAGCGGCGAGAAGATCAAGGGCCTGAACGAGTACATCAACCTGTATAATCAGAAAACCAAGCAGAAGCTGCCTAAGTTTAAGCCACTGTATAAGCAGGTGCTGAGCGATCGGGAGTCTCTGAGCTTCTACGGCGAGGGCTATACATCCGATGAGGAGGTGCTGGAGGTGTTTAGAAACACCCTGAACAAGAACAGCGAGATCTTCAGCTCCATCAAGAAGCTGGAGAAGCTGTTCAAGAATTTTGACGAGTACTCTAGCGCCGGCATCTTTGTGAAGAACGGCCCCGCCATCAGCACAATCTCCAAGGATATCTTCGGCGAGTGGAACGTGATCCGGGACAAGTGGAATGCCGAGTATGACGATATCCACCTGAAGAAGAAGGCCGTGGTGACCGAGAAGTACGAGGACGATCGGAGAAAGTCCTTCAAGAAGATCGGCTCCTTTTCTCTGGAGCAGCTGCAGGAGTACGCCGACGCCGATCTGTCTGTGGTGGAGAAGCTGAAGGAGATCATCATCCAGAAGGTGGATGAGATCTACAAGGTGTATGGCTCCTCTGAGAAGCTGTTCGACGCCGATTTTGTGCTGGAGAAGAGCCTGAAGAAGAACGACGCCGTGGTGGCCATCATGAAGGACCTGCTGGATTCTGTGAAGAGCTTCGAGAATTACATCAAGGCCTTCTTTGGCGAGGGCAAGGAGACAAACAGGGACGAGTCCTTCTATGGCGATTTTGTGCTGGCCTACGACATCCTGCTGAAGGTGGACCACATCTACGATGCCATCCGCAATTATGTGACCCAGAAGCCCTACTCTAAGGATAAGTTCAAGCTGTATTTTCAGAACCCTCAGTTCATGGGCGGCTGGGACAAGGATAAGGAGACAGACTATCGGGCCACCATCCTGAGATACGGCTCCAAGTACTATCTGGCCATCATGGATAAGAAGTACGCCAAGTGCCTGCAGAAGATCGACAAGGACGATGTGAACGGCAATTACGAGAAGATCAACTATAAGCTGCTGCCCGGCCCTAATAAGATGCTGCCAAAGGTGTTCTTTTCTAAGAAGTGGATGGCCTACTATAACCCCAGCGAGGACATCCAGAAGATCTACAAGAATGGCACATTCAAGAAGGGCGATATGTTTAACCTGAATGACTGTCACAAGCTGATCGACTTCTTTAAGGATAGCATCTCCCGGTATCCAAAGTGGTCCAATGCCTACGATTTCAACTTTTCTGAGACAGAGAAGTATAAGGACATCGCCGGCTTTTACAGAGAGGTGGAGGAGCAGGGCTATAAGGTGAGCTTCGAGTCTGCCAGCAAGAAGGAGGTGGATAAGCTGGTGGAGGAGGGCAAGCTGTATATGTTCCAGATCTATAACAAGGACTTTTCCGATAAGTCTCACGGCACACCCAATCTGCACACCATGTACTTCAAGCTGCTGTTTGACGAGAACAATCACGGACAGATCAGGCTGAGCGGAGGAGCAGAGCTGTTCATGAGGCGCGCCTCCCTGAAGAAGGAGGAGCTGGTGGTGCACCCAGCCAACTCCCCTATCGCCAACAAGAATCCAGATAATCCCAAGAAAACCACAACCCTGTCCTACGACGTGTATAAGGATAAGAGGTTTTCTGAGGACCAGTACGAGCTGCACATCCCAATCGCCATCAATAAGTGCCCCAAGAACATCTTCAAGATCAATACAGAGGTGCGCGTGCTGCTGAAGCACGACGATAACCCCTATGTGATCGGCATCGATAGGGGCGAGCGCAATCTGCTGTATATCGTGGTGGTGGACGGCAAGGGCAACATCGTGGAGCAGTATTCCCTGAACGAGATCATCAACAACTTCAACGGCATCAGGATCAAGACAGATTACCACTCTCTGCTGGACAAGAAGGAGAAGGAGAGGTTCGA GGCCCGCCAGAACTGGACCTCCATCGAGAATATCAAGGAGCTGAAGGCCGGCTATATCTCTCAGGTGGTGCACAAGATCTGCGAGCTGGTGGAGAAGTACGATGCCGTGATCGCCCTGGAGGACCTGAACTCTGGCTTTAAGAATAGCCGCGTGAAGGTGGAGAAGCAGGTGTATCAGAAGTTCGAGAAGATGCTGATCGATAAGCTGAACTACATGGTGGACAAGAAGTCTAATCCTTGTGCAACAGGCGGCGCCCTGAAGGGCTATCAGATCACCAATAAGTTCGAGAGCTTTAAGTCCATGTCTACCCAGAACGGCTTCATCTTTTACATCCCTGCCTGGCTGACATCCAAGATCGATCCATCTACCGGCTTTGTGAACCTGCTGAAAACCAAGTATACCAGCATCGCCGATTCCAAGAAGTTCATCAGCTCCTTTGACAGGATCATGTACGTGCCCGAGGAGGATCTGTTCGAGTTTGCCCTGGACTATAAGAACTTCTCTCGCACAGACGCCGATTACATCAAGAAGTGGAAGCTGTACTCCTACGGCAACCGGATCAGAATCTTCCGGAATCCTAAGAAGAACAACGTGTTCGACTGGGAGGAGGTGTGCCTGACCAGCGCCTATAAGGAGCTGTTCAACAAGTACGGCATCAATTATCAGCAGGGCGATATCAGAGCCCTGCTGTGCGAGCAGTCCGACAAGGCCTTCTACTCTAGCTTTATGGCCCTGATGAGCCTGATGCTGCAGATGCGGAACAGCATCACAGGCCGCACCGACGTGGATTTTCTGATCAGCCCTGTGAAGAACTCCGACGGCATCTTCTACGATAGCCGGAACTATGAGGCCCAGGAGAATGCCATCCTGCCAAAGAACGCCGACGCCAATGGCGCCTATAACATCGCCAGAAAGGTGCTGTGGGCCATCGGCCAGTTCAAGAAGGCCGAGGACGAGAAGCTGGATAAGGTGAAGATCGCCATCTCTAACAAGGAGTGGCTGGAGTACGCCCAGACCAGCGTGAAGCACAAAAGGCCGGCGGCCACGAAAAAGGCCGGCCAGGCAAAAAAGAAAAAGGGATCCTACCCATACGATGTTCCAGATTACGCTTATCCCTACGACGTGCCTGATTATGCATACCCATACGATGTCCCCGACTATGCCCTCGAGAGCACCGGTGGCAGCGGAGCTACTAACTTCAGCCTGCTGAAGCAGGCTGGAGACGTGGAGGAGAACCCTGGACCTGCCGGTATGGTGAGCAAGGGCGAGGAGGATAACATGGCCATCATCAAGGAGTTCATGCGCTTCAAGGTGCACATGGAGGGCTCCGTGAACGGCCACGAGTTCGAGATCGAGGGCGAGGGCGAGGGCCGCCCCTACGAGGGCACCCAGACCGCCAAGCTGAAGGTGACCAAGGGTGGCCCCCTGCCCTTCGCCTGGGACATCCTGTCCCCTCAGTTCATGTACGGCTCCAAGGCCTACGTGAAGCACCCCGCCGACATCCCCGACTACTTGAAGCTGTCCTTCCCCGAGGGCTTCAAGTGGGAGCGCGTGATGAACTTCGAGGACGGCGGCGTGGTGACCGTGACCCAGGACTCCTCCCTGCAGGACGGCGAGTTCATCTACAAGGTGAAGCTGCGCGGCACCAACTTCCCCTCCGACGGCCCCGTAATGCAGAAGAAGACCATGGGCTGGGAGGCCTCCTCCGAGCGGATGTACCCCGAGGACGGCGCCCTGAAGGGCGAGATCAAGCAGAGGCTGAAGCTGAAGGACGGCGGCCACTACGACGCTGAGGTCAAGACCACCTACAAGGCCAAGAAGCCCGTGCAGCTGCCCGGCGCCTAC AACGTCAACATCAAGTTGGACATCACCTCCCACAACGAGGACTACACCATCGTGGAACAGTACGAACGCGC CGAGGGCCGCCACTCCACCGGCGGCATGGACGAGCTGTACAAGTAA

#### • Expression of the SpCas9 nuclease

Green: 3XFlag, Orange: SV40 NLS, blue: SpCas9, violet: nucleoplasmin NLS, yellow: P2A, Red: mCherry GACTACAAAGACCATGACGGTGATTATAAAGATCATGACATCGATTACAAGGATGACGATGACAAGATGGCCC CCAAGAAGAAGAGGAAGGTGGGCATTCACCGCGGGGTACCCATGGACAAGAAGTACTCCATTGGGCTCGAT ATCGGCACAAACAGCGTCGGCTGGGCCGTCATTACGGACGAGTACAAGGTGCCGAGCAAAAAATTCAAAGTTCTGGGCAATACCGATCGCCACAGCATAAAGAAGAACCTCATTGGCGCCCTCCTGTTCGACTCCGGGGAGACGGCCGAAGCCACGCGGCTCAAAAGAACAGCACGGCGCAGATATACCCGCAGAAAGAATCGGATCTGCTACCTGCAGGAGATCTTTAGTAATGAGATGGCTAAGGTGGATGACTCTTTCTTCCATAGGCTGGAGGAGTCCTTTTTGGTGGAGGAGGATAAAAAGCACGAGCGCCACCCAATCTTTGGCAATATCGTGGACGAGGTGGCGTACCATGAAAAGTACCCAACCATATATCATCTGAGGAAGAAGCTTGTAGACAGTACTGATAAGGCTGACTTGCGGTTGATCTATCTCGCGCTGGCGCATATGATCAAATTTCGGGGACACTTCCTCATCGAGGGGGACCTGAACCCAGACAACAGCGATGTCGACAAACTCTTTATCCAACTGGTTCAGACTTACAATCAGCTTTTCGAAGAGAACCCGATCAACGCATCCGGAGTTGACGCCAAAGCAATCCTGAGCGCTAGGCTGTCCAAATCCCGGCGGCTCGAAAACCTCATCGCACAGCTCCCTGGGGAGAAGAAGAACGGCCTGTTTGGTAATCTTATCGCCCTGTCACTCGGGCTGACCCCCAACTTTAAATCTAACTTCGACCTGGCCGAAGATGCCAAGCTTCAACTGAGCAAAGACACCTACGATGATGATCTCGACAATCTGCTGGCCCAGATCGGCGACCAGTACGCAGACCTTTTTTTGGCGGCAAAGAACCTGTCAGACGCCATTCTGCTGAGTGATATTCTGCGAGTGAACACGGAGATCACCAAAGCTCCGCTGAGCGCTAGTATGATCAAGCGCTATGATGAGCACCACCAAGACTTGACTTTGCTGAAGGCCCTTGTCAGACAGCAACTGCCTGAGAAGTACAAGGAAATTTTCTTCGATCAGTCTAAAAATGGCTACGCCGGATACATTGACGGCGGAGCAAGCCAGGAGGAATTTTACAAATTTATTAAGCCCATCTTGGAAAAAATGGACGGCACCGAGGAGCTGCTGGTAAAGCT TAACAGAGAAGATCTGTTGCGCAAACAGCGCACTTTCGACAATGGAAGCATCCCCCACCAGATTCACCTGGGCGAACTGCACGCTATCCTCAGGCGGCAAGAGGATTTCTACCCCTTTTTGAAAGATAACAGGGAAAAGATTGAGAAAATCCTCACATTTCGGATACCCTACTATGTAGGCCCCCTCGCCCGGGGAAATTCCAGATTCGCGTGGATGACTCGCAAATCAGAAGAGACCATCACTCCCTGGAACTTCGAGGAAGTCGTGGATAAGGGGGCCTCTGCCCAGTCCTTCATCGAAAGGATGACTAACTTTGATAAAAATCTGCCTAACGAAAAGGTGCTTCCTAAACACTCTCTGCTGTACGAGTACTTCACAGTTTATAACGAGCTCACCAAGGTCAAATACGTCACAGAAGGGATGAGAAAGCCAGCATTCCTGTCTGGAGAGCAGAAGAAAGCTATCGTGGACCTCCTCTTCAAGACGAACCGGAAAGTTACCGTGAAACAGCTCAAAGAAGACTATTTCAAAAAGATTGAATGTTTCGACTCTGTTGAAATCAGCGGAGTGGAGGATCGCTTCAACGCATCCCTGGGAACGTATCACGATCTCCTGAAAATCATTAAAGACAAGGACTTCCTGGACAATGAGGAGAACGAGGACATTCTTGAGGACATTGTCCTCACCCTTACGTTGTTTGAAGATAGGGAGATGATTGAAGAACGCTTGAAAACTTACGCTCATCTCTTCGACGACAAAGTCATGAAACAGCTCAAGAGGCGCCGATATACAGGATGGGGGCGGCTGTCAAGAAAACTGATCAATGGGATCCGAGACAAGCAGAGTGGAAAGACAATCCTGGATTTTCTTAAGTCCGATGGATTTGCCAACCGGAACTTCATGCAGTTGATCCATGATGACTCTCTCACCTTTAAGGAGGACATCCAGAAAGCACAAGTTTCTGGCCAGGGGGACAGTCTTCACGAGCACATCGCTAATCTTGCAGGTAGCCCAGCTATCAAAAAGGGAATACTGCAGACCGTTAAGGTCGTGGATGAACTCGTCAAAGTAATGGGAAGGCATAAGCCCGAGAATATCGTTATCGAGATGGCCCGAGAGAACCAAACTACCCAGAAGGGACAGAAGAACAGTAGGGAAAGGATGAAGAGGATTGAAGAGGGTATAAAAGAACTGGGGTCCCAAATCCTTAAGGAACACCCAGTTGAAAACACCCAGCTTCAGAATGAGAAGCTCTACCTGTACTACCTGCAGAACGGCAGGGACATGTACGTGGATCAGGAACTGGACATCAATCGGCTCTCCGACTACGACGTGGATCATATCGTGCCCCAGTCTTTTCTCAAAGATGATTCTATTGATAATAAAGTGTTGACAAGATCCGATAAAAATAGAGGGAAGAGTGATAACGTCCCCTCAGAAGAAGTTGTCAAGAAAATGAAAAATTATTGGCGGCAGCTGCTGAACGCCAAACTGATCACACAACGGAAGTTCGATAATCTGACTAAGGCTGAACGAGGTGGCCTGTCTGAGTTGGATAAAGCCGGCTTCATCAAAAGGCAGCTTGTTGAGACACGCCAGATCACCAAGCACGTGGCCCAAATTCTCGATTCACGCATGAACACCAAGTACGATGAAAATGACAAACTGATTCGAGAGGTGAAAGTTATTACTCTGAAGTCTAAGCTGGTCTCAGATTTCAGAAAGGACTTTCAGTTTTATAAGGTGAGAGAGATCAACAATTACCACCATGCGCATGATGCCTACCTGAATGCAGTGGTAGGCACTGCACTTATCAAAAAATATCCCAAGCTTGAATCTGAATTTGTTTACGGAGACTATAAAGTGTACGATGTTAGGAAAATGATCGCAAAGTCTGAGCAGGAAATAGGCAAGGCCACCGCTAAGTACTTCTTTTACAGCAATATTATGAATTTTTTCAAGACCGAGATTACACTGGCCAATGGAGAGATTCGGAAGCGACCACTTATCGAAACAAACGGAGAAACAGGAGAAATCGTGTGGGACAAGGGTAGGGATTTCGCGACAGTCCGGAAGGTCCTGTCCATGCCGCAGGTGAACATCGTTAAAAAGACCGAAGTACAGACCGGAGGCTTCTCCAAGGAAAGTATCCTCCCGAAAAGGAACAGCGACAAGCTGATCGCACGCAAAAAAGATTGGGACCCCAAGAAATACGGCGGATTCGATTCTCCTACAGTCGCTTACAGTGTACTGGTTGTGGCCAAAGTGGAGAAAGGGAAGTCTAAAAAACTCAAAAGCGTCAAGGAACTGCTGGGCATCACAATCATGGAGCGATCAAGCTTCGAAAAAAACCCCATCGACTTTCTCGAGGCGAAAGGATATAAAGAGGTCAAAAAAGACCTCATCATTAAGCTTCCCAAGTACTCTCTCTTTGAGCTTGAAAACGGCCGGAAACGAATGCTCGCTAGTGCGGGCGAGCTGCAGAAAGGTAACGAGCTGGCACTGCCCTCTAAATACGTTAATTTCTTGTATCTGGCCAGCCACTATGAAAAGCTCAAAGGGTCTCCCGAAGATAATGAGCAGAAGCAGCTGTTCGTGGAACAACACAAACACTACCTTGATGAGATCATCGAGCAAATAAGCGAATTCTCCAAAAGAGTGATCCTCGCCGACGCTAACCTCGATAAGGTGCTTTCTGCTTACAATAAGCACAGGGATAAGCCCATCAGGGAGCAGGCAGAAAACATTATCCACTTGTTTACTCTGACCAACTTGGGCGCGCCTGCAGCCTTCAAGTACTTCGACACCACCATAGACAGAAAGCGGTACACCTCTACAAAGGAGGTCCTGGACGCCACACTGATTCATCAGTCAATTACGGGGCTCTATGAAACAAGAATCGACCTCTCTCAGCTCGGTGGAGACAGCAGGGCTGACCCCAAGAAGAAGAGGAAGGTGACCGGTGGCAGCGGAGCTACTAACTTCAGCCTGCTGAAGCAGGCTGGAGACGTGGAGGAGAACCCTGGACCTGCCGGTGCCGGTATGGTGAGCAAGGGCGAGGAGGATAACATGGCCATCATCAAGGAGTTCATGCGCTTCAAGGTGCACATGGAGGGCTCCGTGAACGGCCACGAGTTCGAGATCGAGGGCGAGGGCGAGGGCCGCCCCTACGAGGGCACCCAGACCGCCAAGCTGAAGGTGACCAAGGGTGGCCCCCTGCCCTTCGCCTGGGACATCCTGTCCCCTCAGTTCATGTACGGCTCCAAGGCCTACGTGAAGCACCCCGCCGACATCCCCGACTACTTGAAGCTGTCCTTCCCCGAGGGCTTCAAGTGGGAGCGCGTGATGAACTTCGAGGACGGCGGCGT GGTGACCGTGACCCAGGACTCCTCCCTGCAGGACGGCGAGTTCATCTACAAGGTGAAGCTGCGCGGCACC AACTTCCCCTCCGACGGCCCCGTAATGCAGAAGAAGACCATGGGCTGGGAGGCCTCCTCCGAGCGGATGT ACCCCGAGGACGGCGCCCTGAAGGGCGAGATCAAGCAGAGGCTGAAGCTGAAGGACGGCGGCCACTACG ACGCTGAGGTCAAGACCACCTACAAGGCCAAGAAGCCCGTGCAGCTGCCCGGCGCCTACAACGTCAACAT CAAGTTGGACATCACCTCCCACAACGAGGACTACACCATCGTGGAACAGTACGAACGCGCCGAGGGCCGC CACTCCACCGGCGGCATGGACGAGCTGTACAAGTAA

**Supplementary Figure 1.**
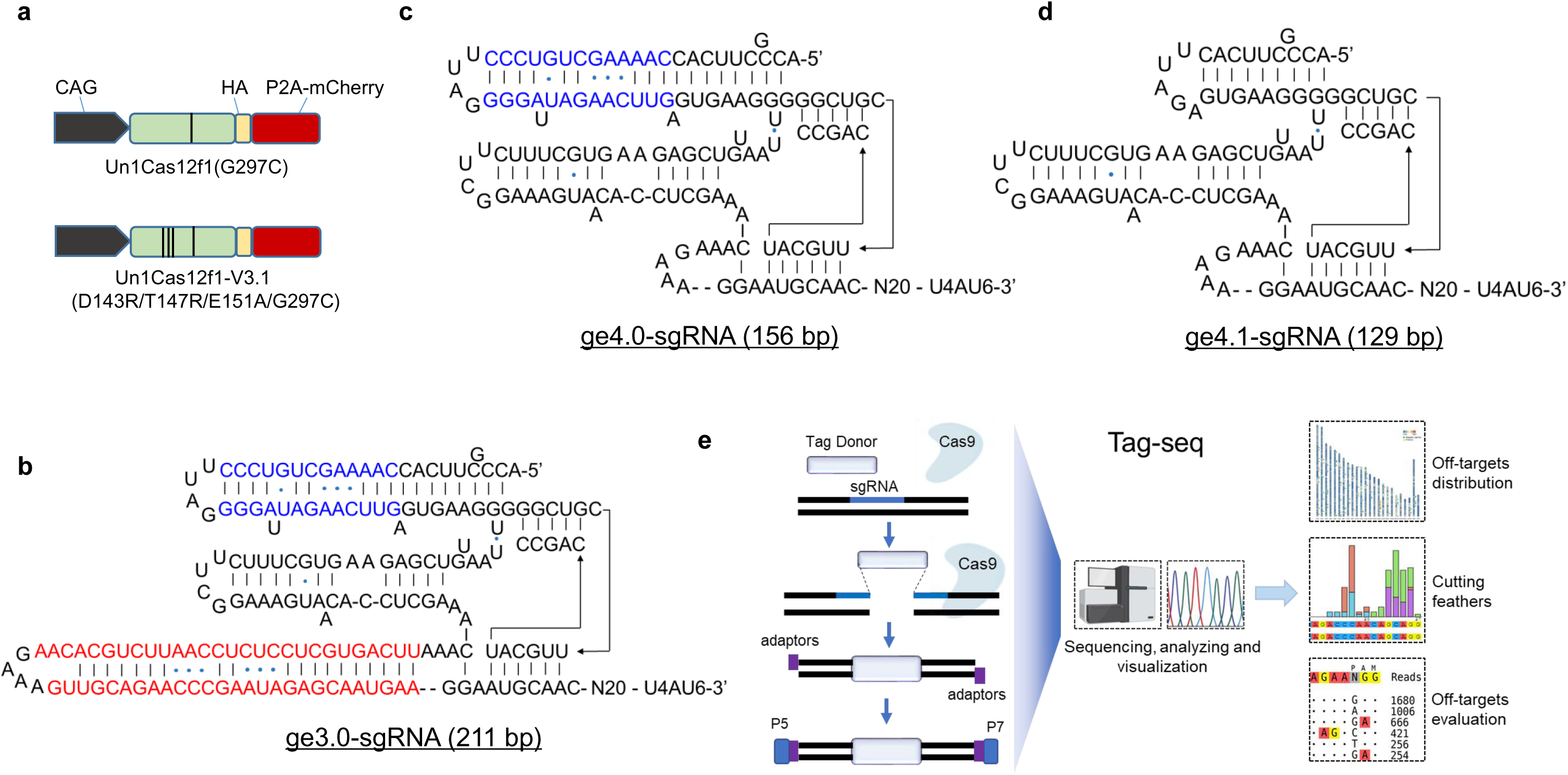
The schematic of the engineered CRISPR-Un1Cas12f1 system and the Tag-seq method. **a** The structures of the Un1Cas12f1 plasmids, where the sequences of the WT Un1Cas12f1-nucleases contained the G297C mutation was from the previous report^8^. **b-d** The structures and detailed sequences of the engineered sgRNAs, in which the colored sequences were the differences among these three sgRNAs. The ge4.0-sgRNA (c) was generated by the deletion of the red sequences in ge3.0-sgRNA (b), and the ge4.1-sgRNA (d) was generated by the deletion of the blue sequences in ge4.0-sgRNA. **e** The workflow of Tag-seq, a streamlined sequencing method, which has a broad spectrum of applications, like tracing DNA double-strand breaks induced by CRISPR tools, profiling CRISPR-based off-targets, evaluation of gene editing events, and profiling molecular characteristics in on- and off-target sites *etc*.

**Supplementary Figure 2.**
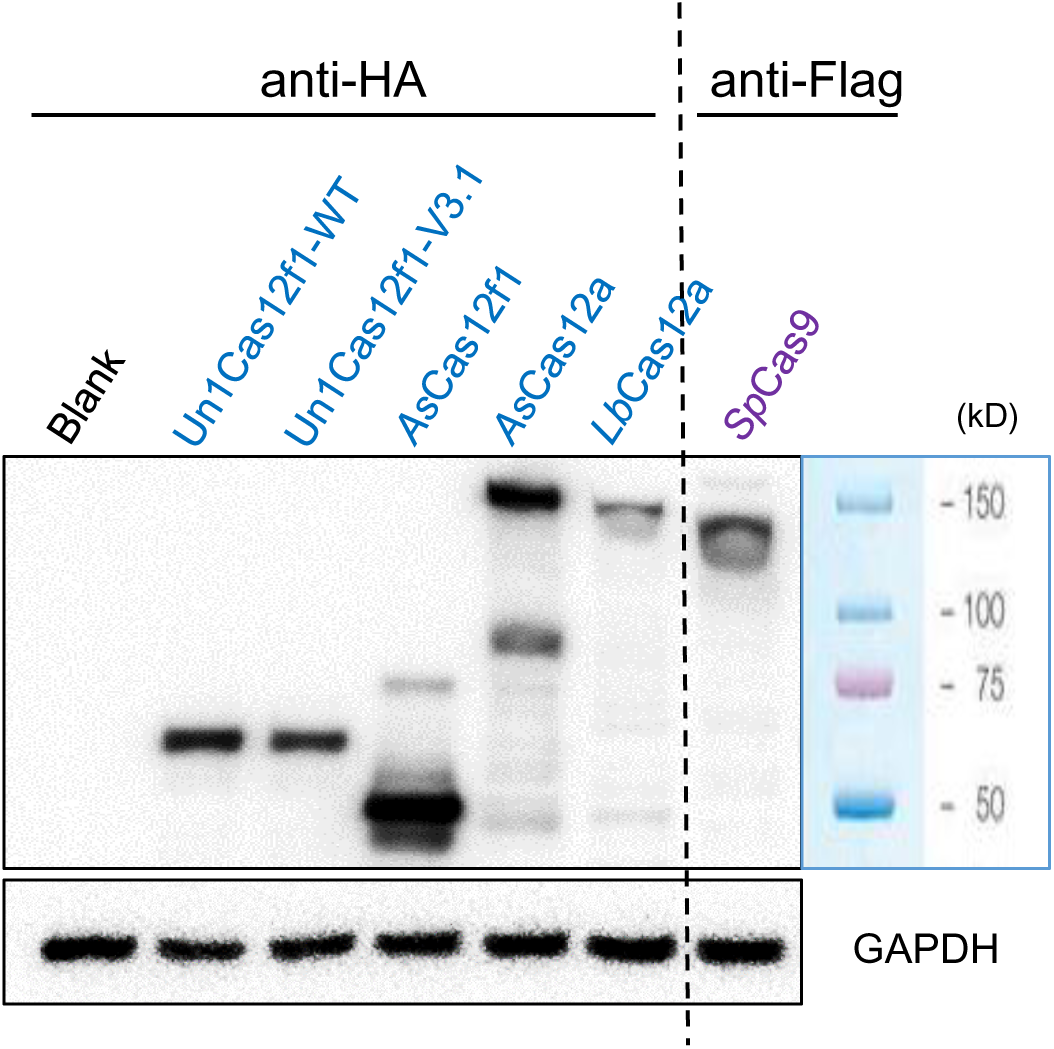
The expression of the Cas nucleases. Western blot showing the expression levels of the Cas-protein nucleases. Exception for the *Sp*Cas9 that fusing with the anti-Flag at the N-terminus, the other nuclease were detected by the anti-HA which fused to the C-terminus. Blank, HEK293T without transfection.

**Supplementary Figure 3.**
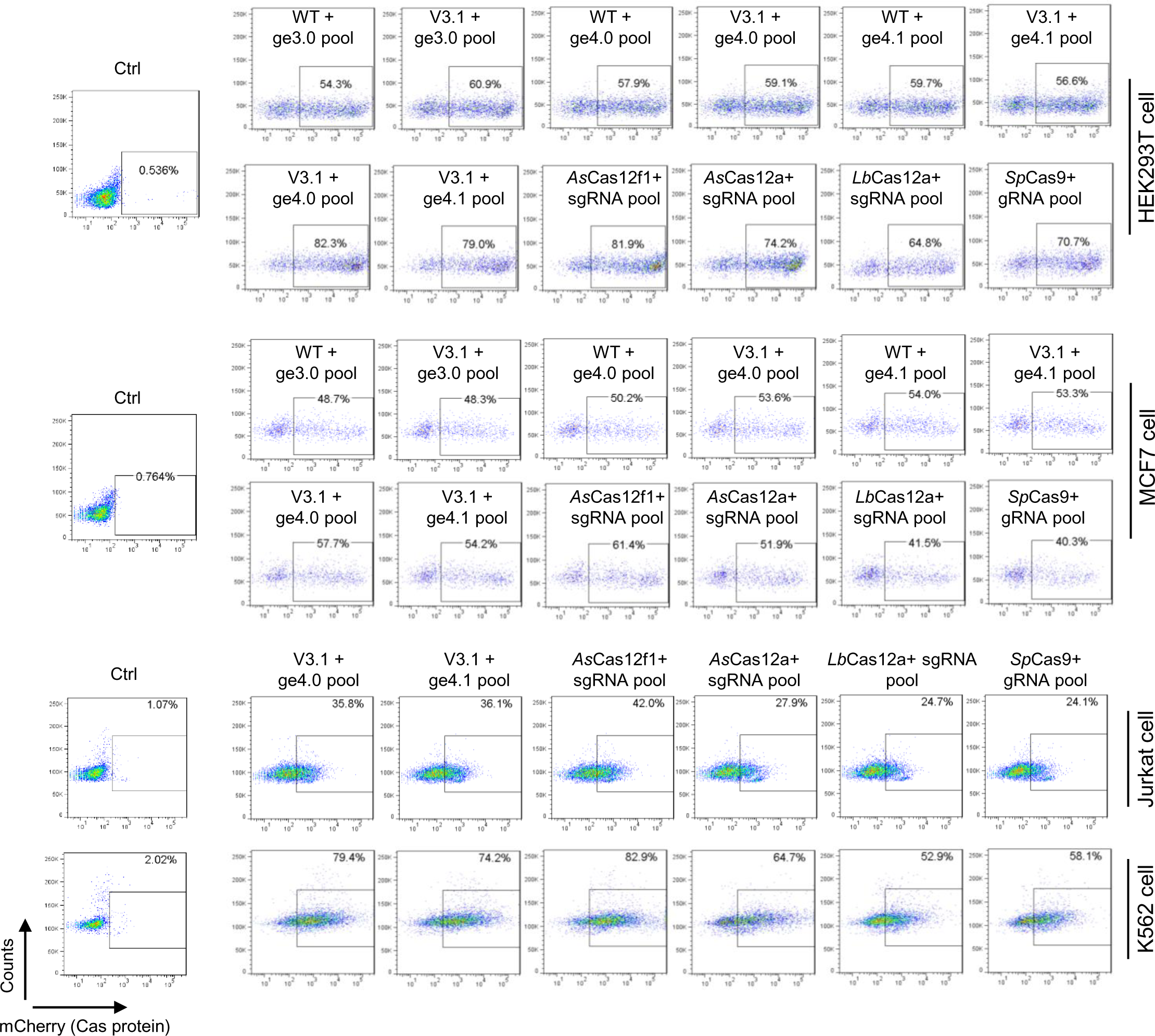
Detection of the transfection efficiency by FACS. The transfection were administrated by PEI-based method (for HEK293T and MCF7 cells) and Lonza kit (2D, for Jurkat and K562 cells) with the plasmids expression of Cas-protein (fusing a P2A mCherry reporter) and the pooled sgRNAs, and an Tag-oligo DNA, and the transfection efficiency was determined by FACS with the mCherry reporter.

**Supplementary Figure 4.**
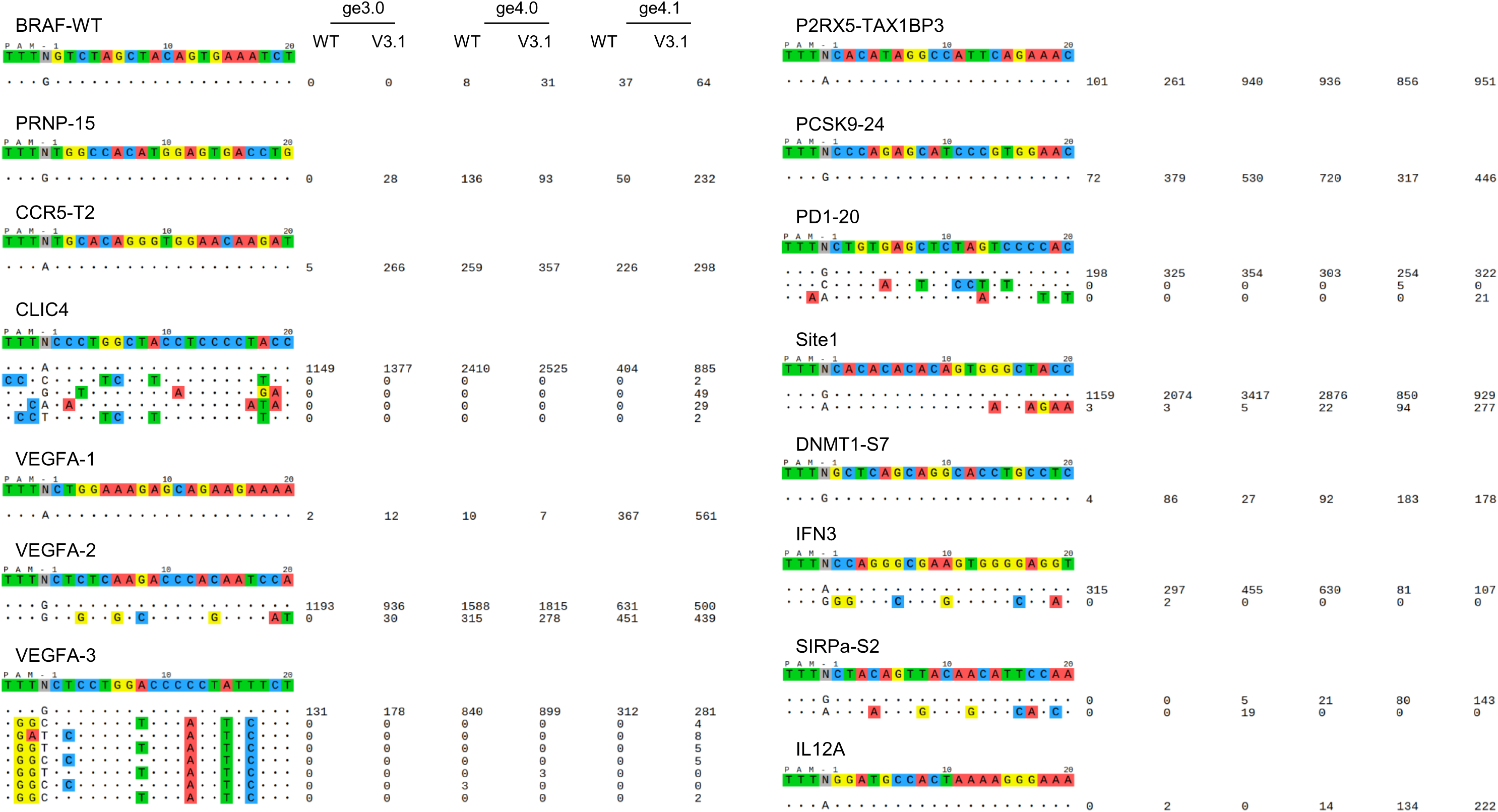
Specificity comparison of the engineered CRISPR-Un1Cas12f1 system by Tag-seq in HEK293T cells. HEK293T cells were transfected by PEI method with the plasmids expressing Un1Cas12f1-WT or -V3.1 and a pooled twenty-one sgRNAs (containing the ge3.0, ge4.0, or ge4.1), and the Tag-oligo DNA sequence. Genomic DNA was harvested three days post-transfection for libraries construction and Tag-seq analysis. Read counts represented a measure of cleavage frequency at a given site, mismatched positions within the spacer or PAM are highlighted in color (also see Fig. 1b).

**Supplementary Figure 6.**
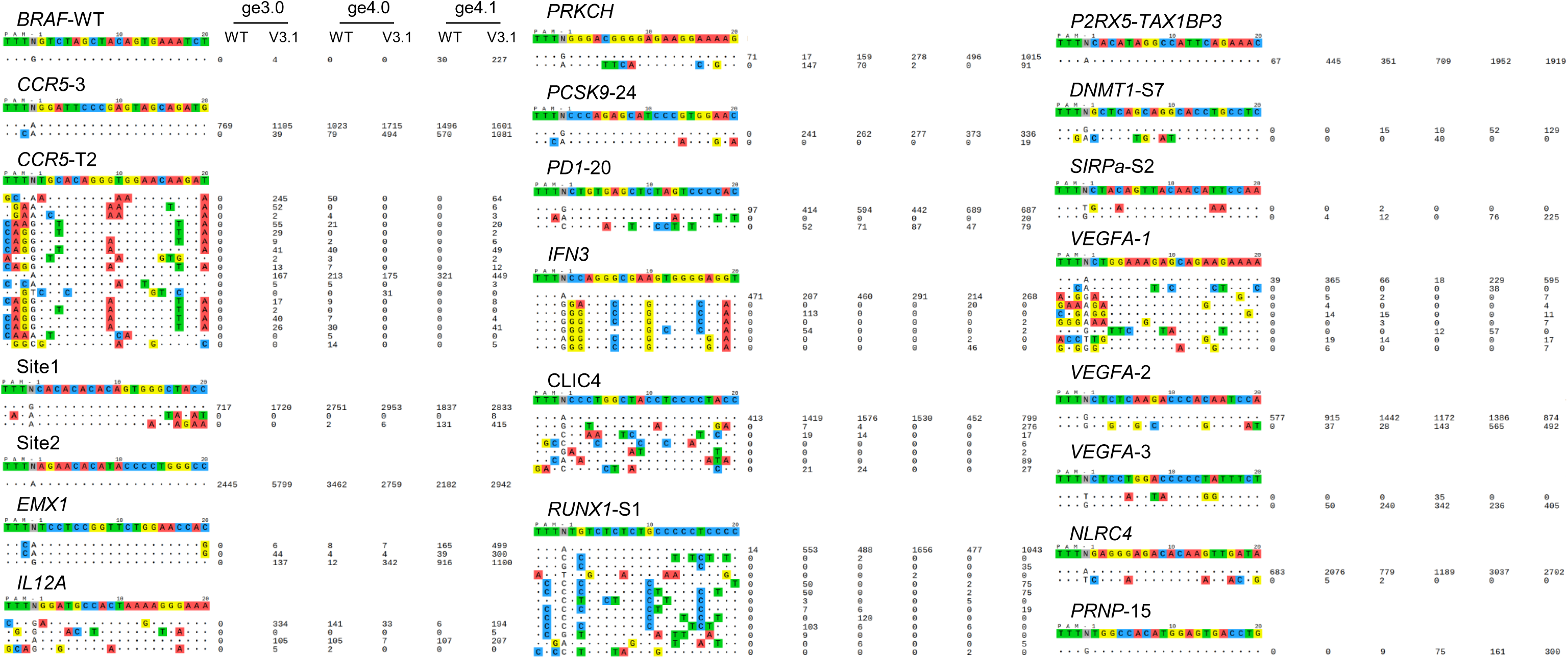
Specificity comparison of the engineered CRISPR-Un1Cas12f1 system by Tag-seq in MCF7 cells. MCF7 cells were transfected by PEI method with the plasmids expressing Un1Cas12f1-WT or -V3.1 and a pooled twenty-one sgRNAs (containing the ge3.0, ge4.0, or ge4.1), and the Tag-oligo DNA sequence. Genomic DNA was harvested three days post-transfection for libraries construction and Tag-seq analysis. Read counts represented a measure of cleavage frequency at a given site, mismatched positions within the spacer or PAM are highlighted in color.

**Supplementary Figure 7.**
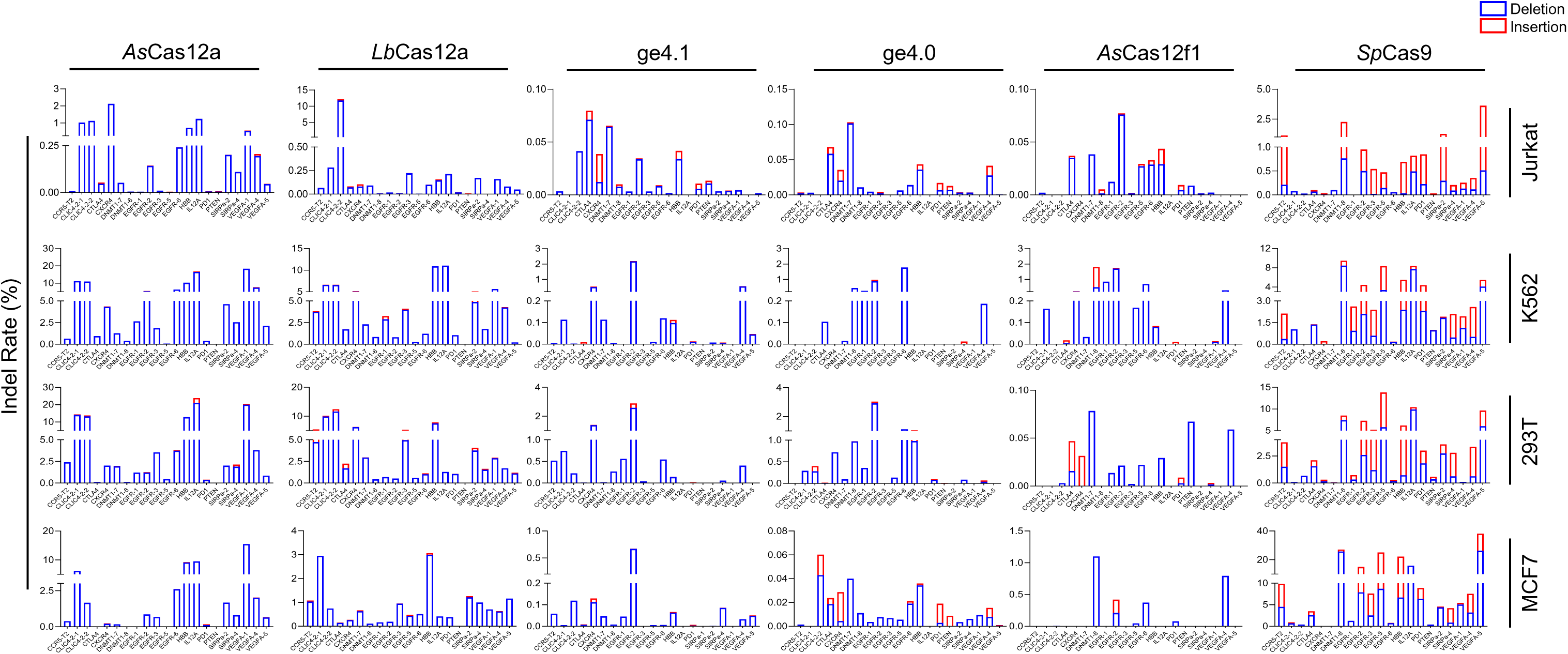
Activity comparison of the DNA editing CRISPR system by Deep-seq in 4 cell lines. Deep-seq revealed the editing activities of the DNA CRISPR editors by targeting twenty-one sites in various cell lines, including HEK293T, MCF7, K562, and Jurkat. Cells were transfected with the plasmids expressing DNA-editing nucleases, the corresponding sgRNAs that pooled with twenty-one guides. Two days post-transfection genomic DNA was harvested for Deep-seq libraries construction and editing efficiency analyses.

**Supplementary Figure 8.**
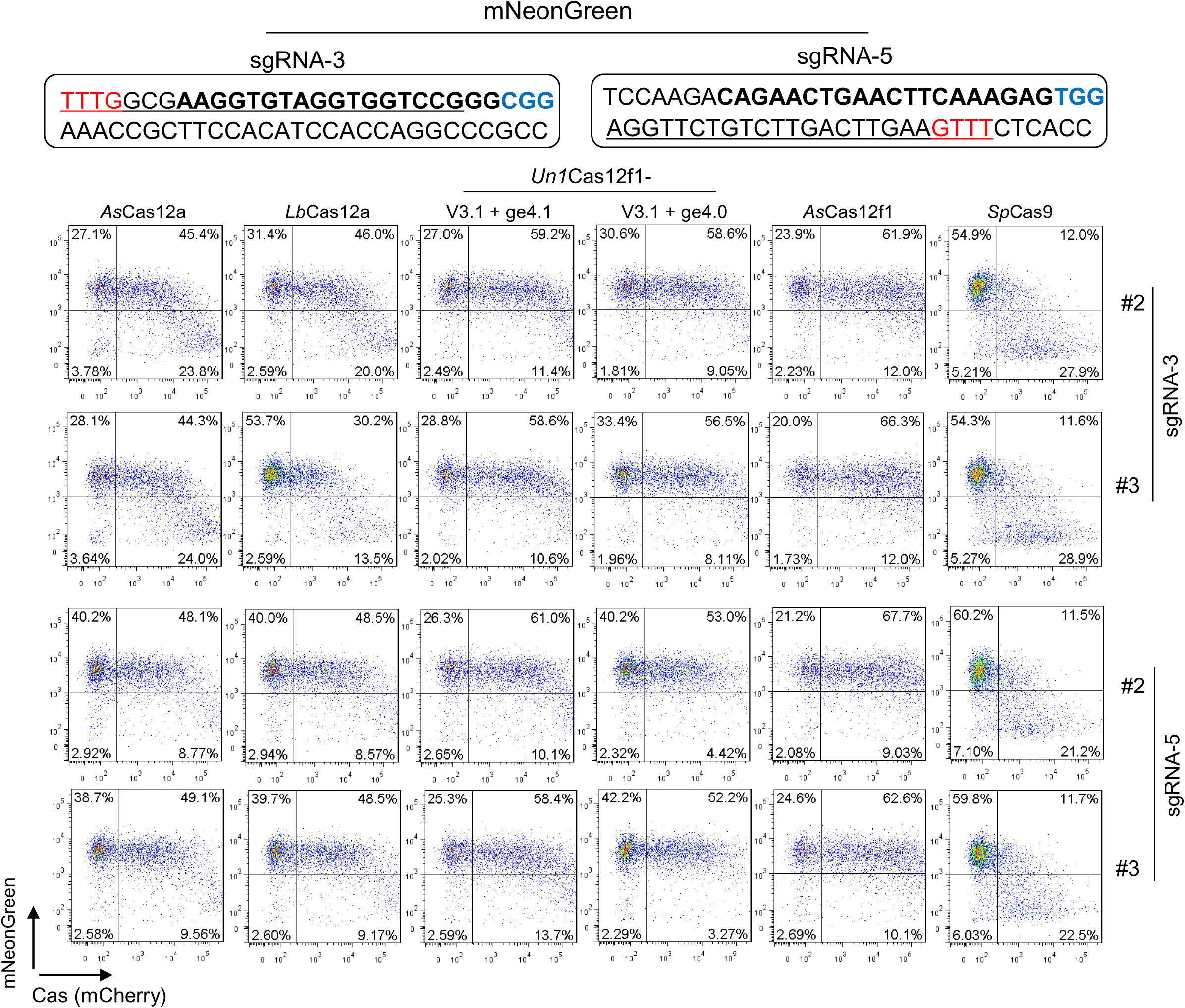
Activity comparison of the DNA editing CRISPR system by FACS in HEK293T-KI-mNeonGreen reporter cells. Another two replicates for the Fig. 3c. The editing efficiency was determined as the proportion of GFP negative cells within the Cas-nucleases transfected cells (mCherry-positive) by FACS. mNeonGree-sgRNA3/5, DNA editing CRISPR targeting mNeonGreen site 3/5. Red sequences showing the PAM of Cas12f/Cas12a, Blue sequences showing the PAM of Cas9. n=3 independent experiments.

**Supplementary Figure 9.**
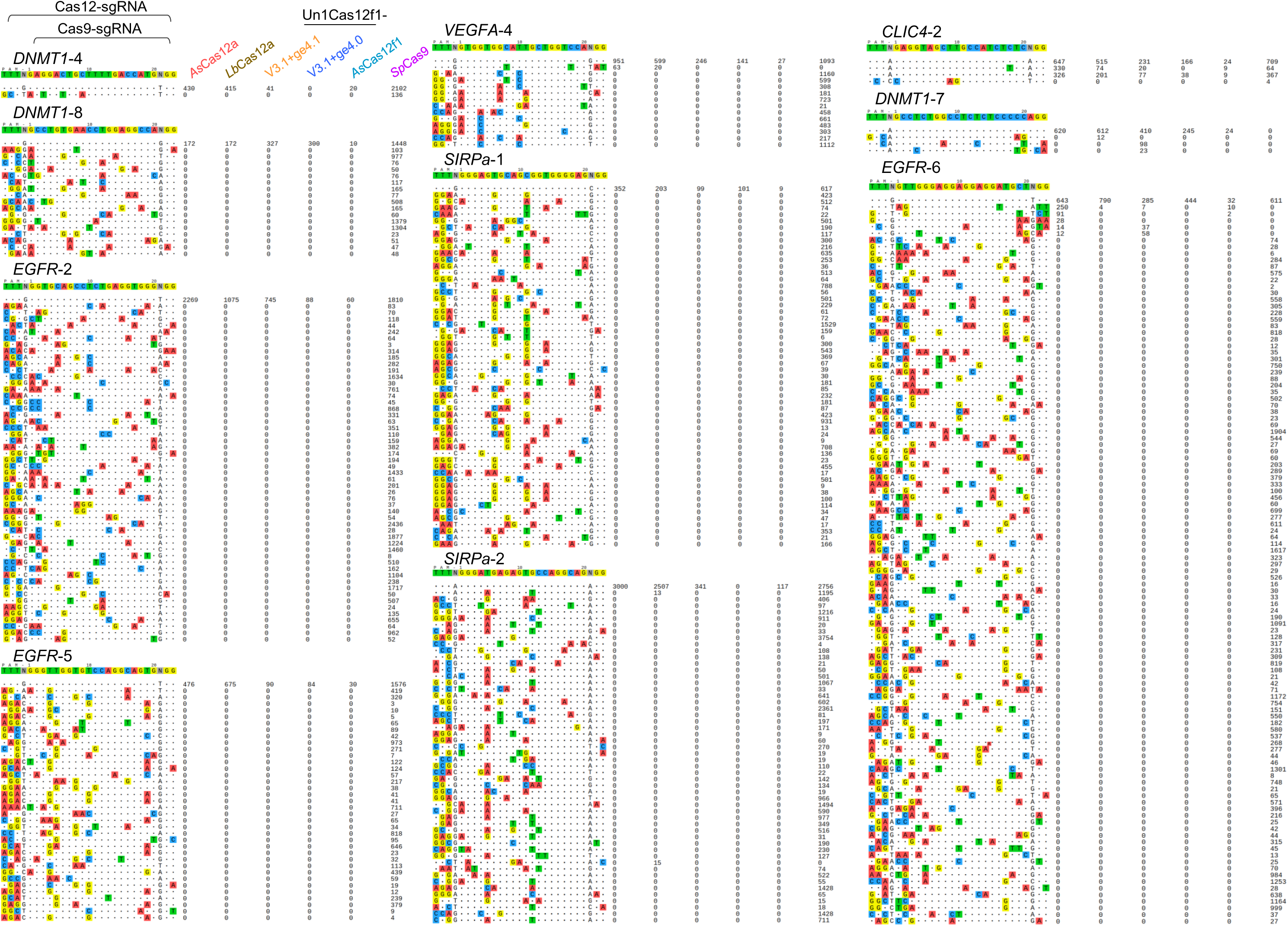
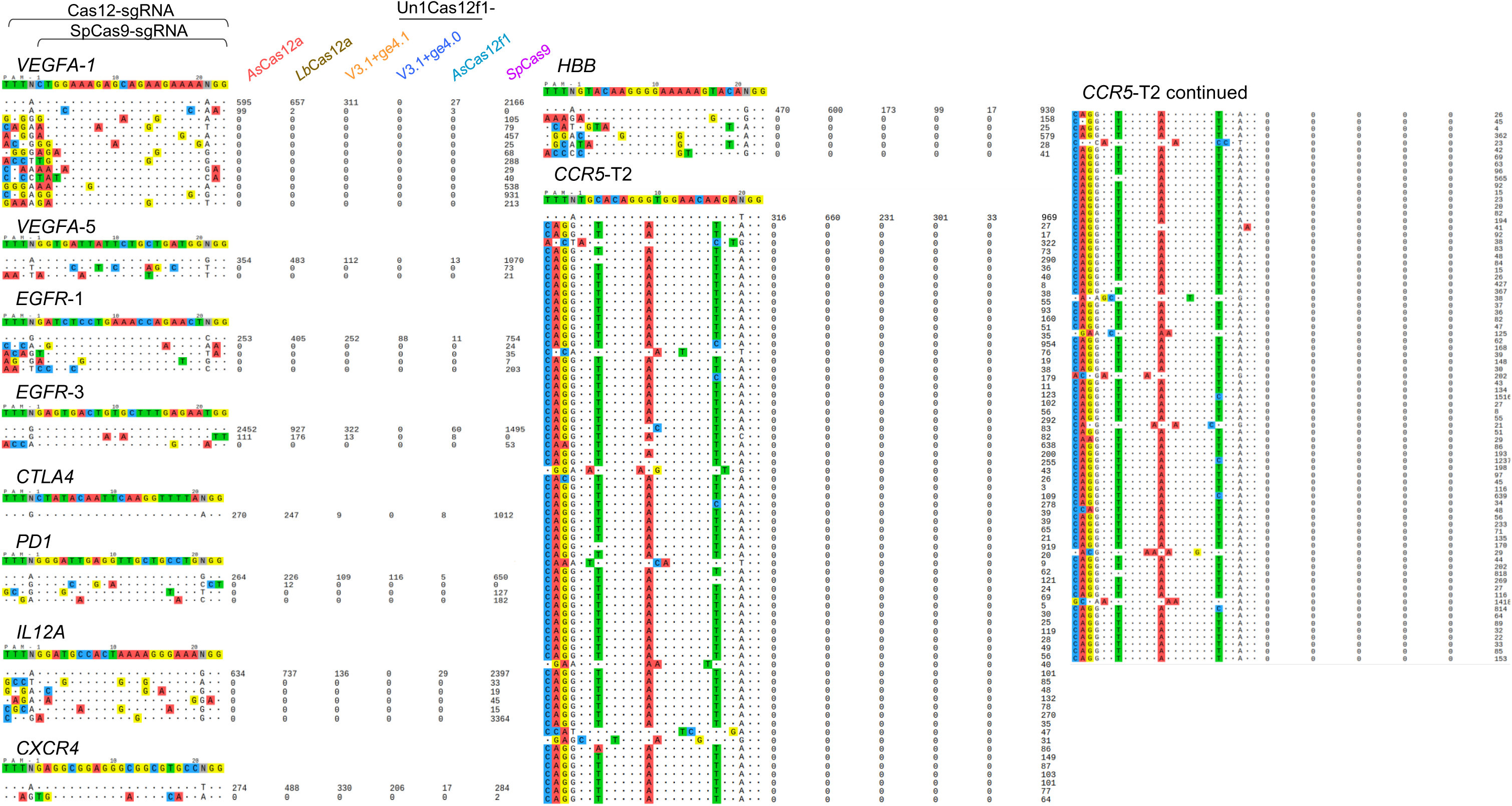
Specificity comparison of the DNA editing CRISPR systems in HEK293T cells. HEK293T cells were transfected by PEI method with the plasmids expressing DNA-editing nucleases, the corresponding sgRNAs that pooled with twenty-one guides, and the Tag-oligo DNA sequence. Genomic DNA was harvested three days post-transfection for libraries construction and Tag-seq analysis. Read counts represented a measure of cleavage frequency at a given site, mismatched positions within the spacer or PAM are highlighted in color (also see Fig. 2b). The targeted sites for Cas12a, Cas12f1 and *Sp*Cas9 share with a common spacer sequence as shown in the top.

**Supplementary Figure 10.**
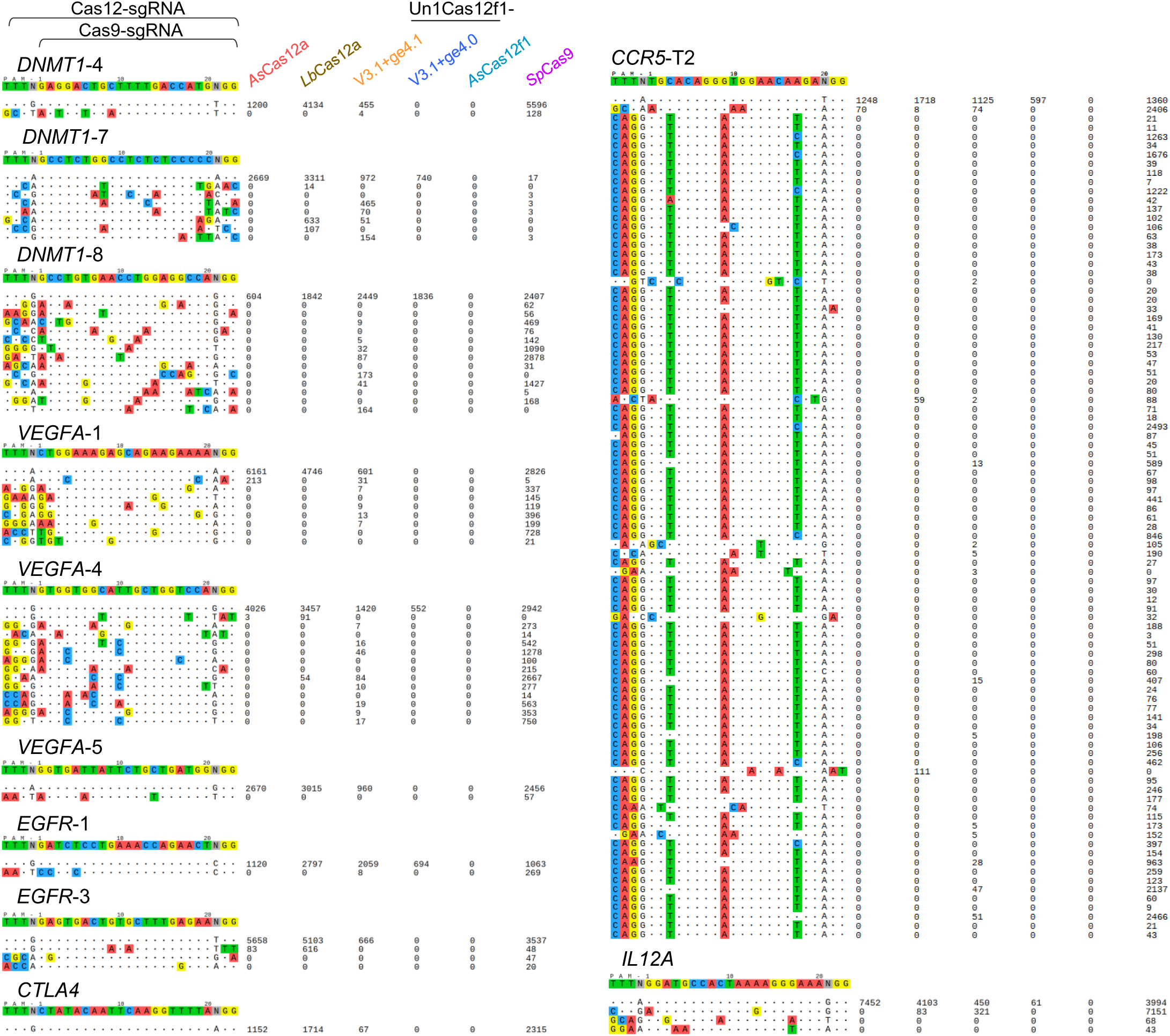
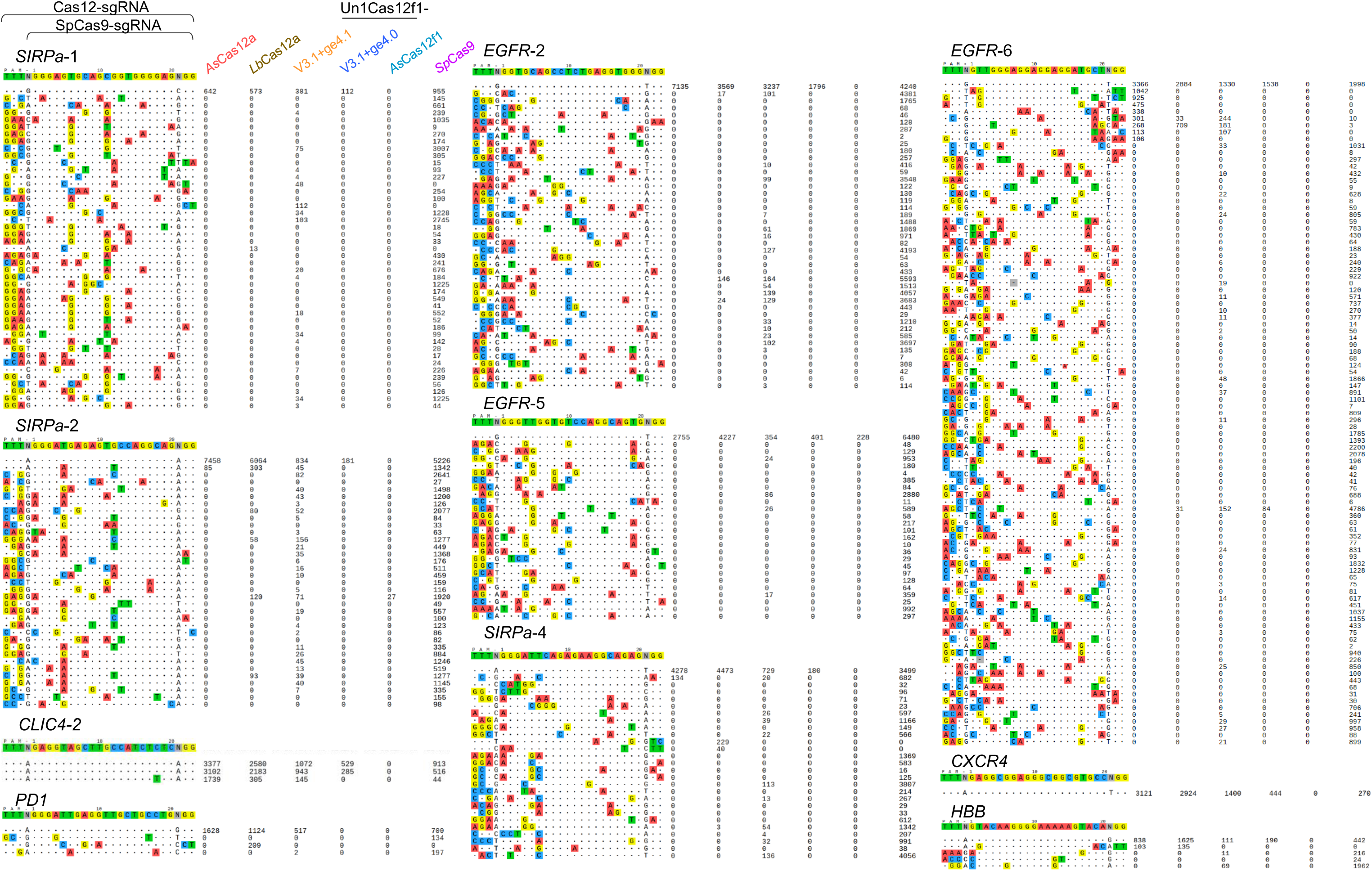
Specificity comparison of the DNA editing CRISPR systems in MCF7 cells. MCF7 cells were transfected by PEI method with the plasmids expressing DNA-editing nucleases, the corresponding sgRNAs that pooled with twenty-one guides, and the Tag-oligo DNA sequence. Genomic DNA was harvested three days post-transfection for libraries construction and Tag-seq analysis. Read counts represented a measure of cleavage frequency at a given site, mismatched positions within the spacer or PAM are highlighted in color. The targeted sites for Cas12a, Cas12f1 and *Sp*Cas9 share with a common spacer sequence as shown in the top.

**Supplementary Figure 11.**
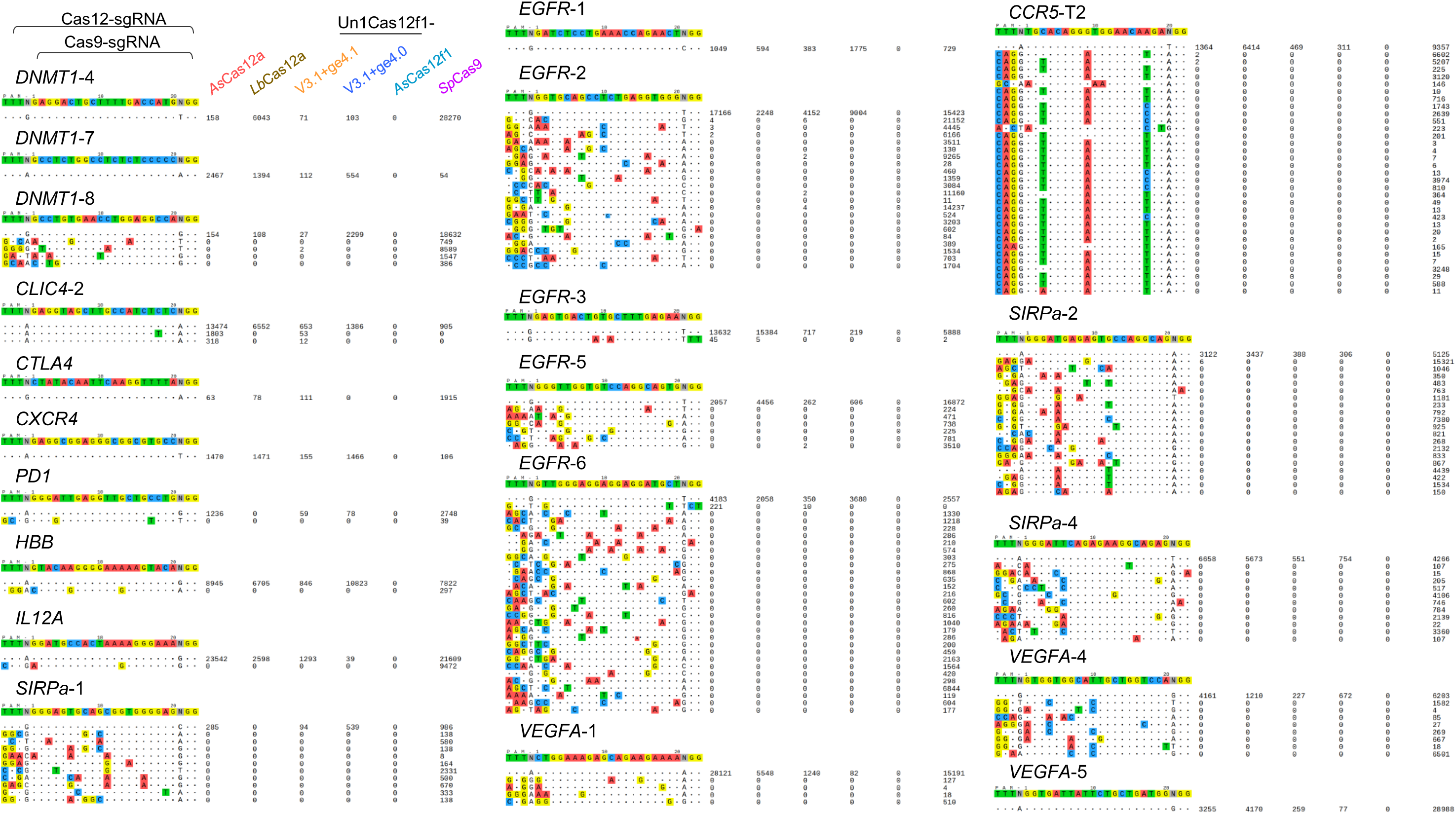
Specificity comparison of the DNA editing CRISPR systems in K562 cells. K562 cells were co-transfected by Lonza electroporation method with the plasmids expressing DNA-editing nucleases, the corresponding sgRNAs that pooled with twenty-one guides, and the Tag-oligo DNA sequence. Genomic DNA was harvested three days post-transfection for libraries construction and Tag-seq analysis. Read counts represented a measure of cleavage frequency at a given site, mismatched positions within the spacer or PAM are highlighted in color. The targeted sites for Cas12a, Cas12f1 and *Sp*Cas9 share a common spacer sequence as shown in the top.

**Supplementary Figure 12.**
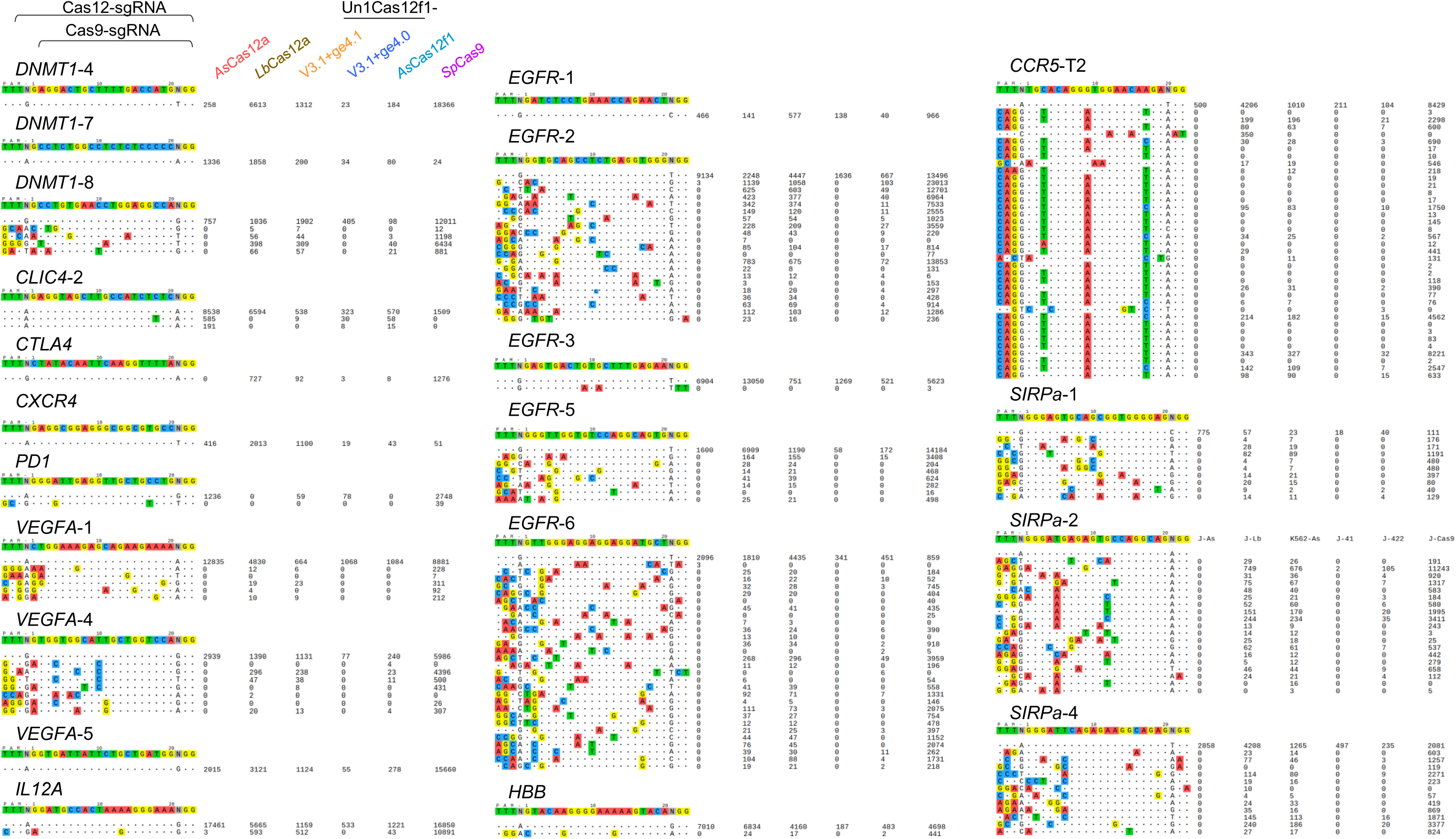
Specificity comparison of the DNA editing CRISPR systems in Jurkat cells. Jurkat cells were transfected by Lonza electroporation method with the plasmids expressing DNA-editing nucleases, the corresponding sgRNAs that pooled with twenty-one guides, and the Tag-oligo DNA sequence. Genomic DNA was harvested three days post-transfection for libraries construction and Tag-seq analysis. Read counts represented a measure of cleavage frequency at a given site, mismatched positions within the spacer or PAM are highlighted in color. The targeted sites for Cas12a, Cas12f1 and *Sp*Cas9 share a common spacer sequence as shown in the top.

**Table S1.**
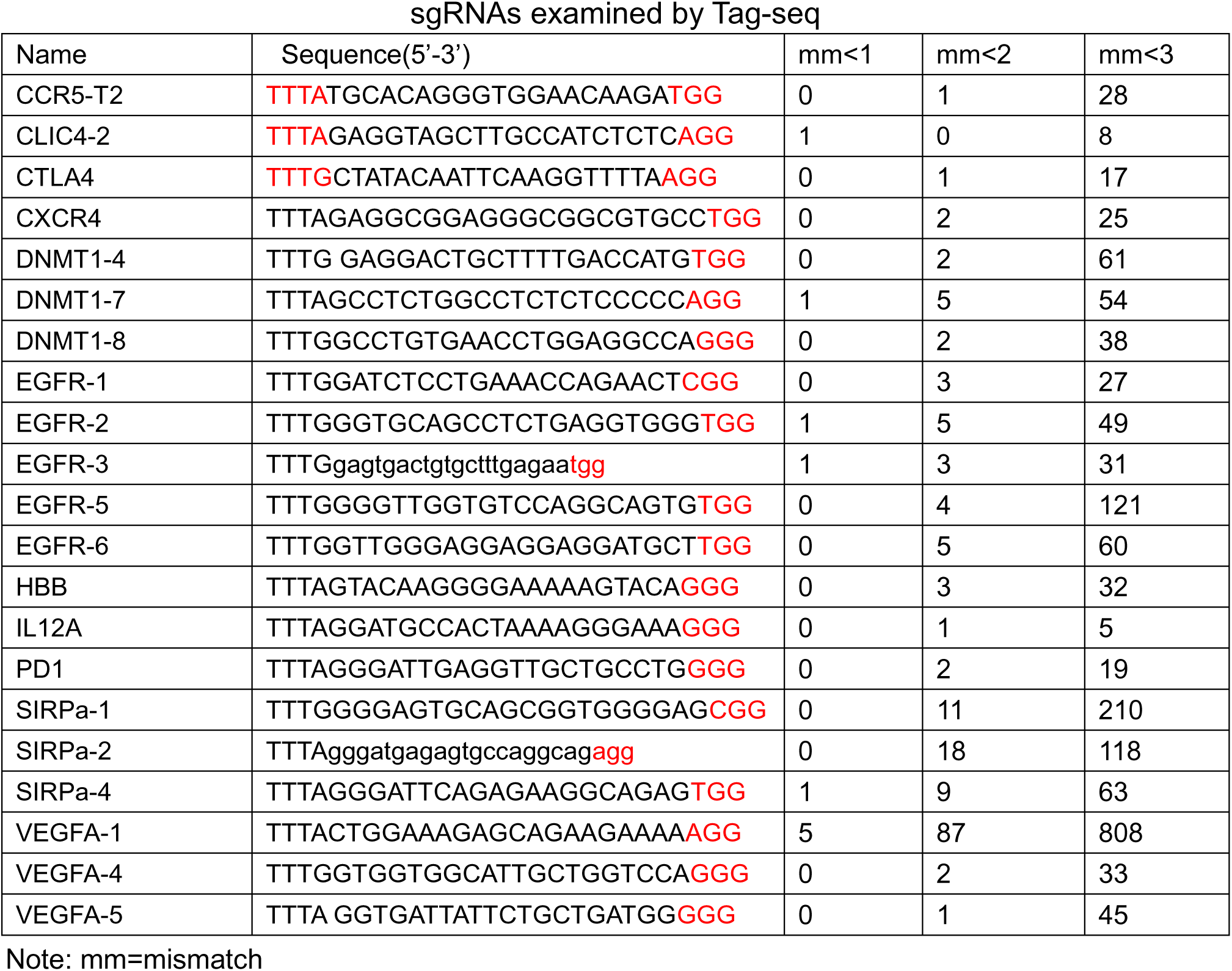
Summary of potential mismatched sites in the reference human genome for the 21 sgRNAs examined by Tag-seq

**Table S2.**
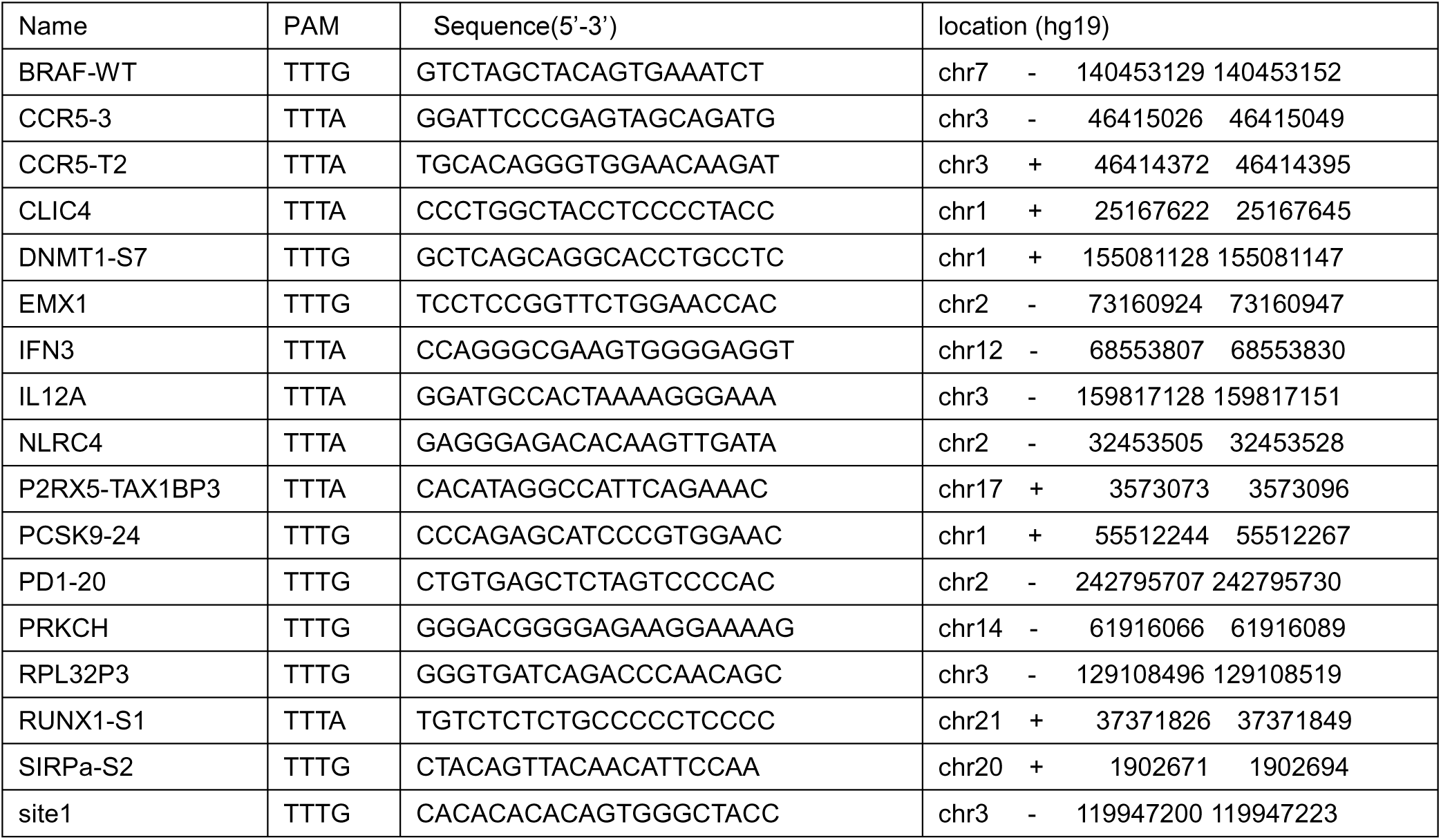
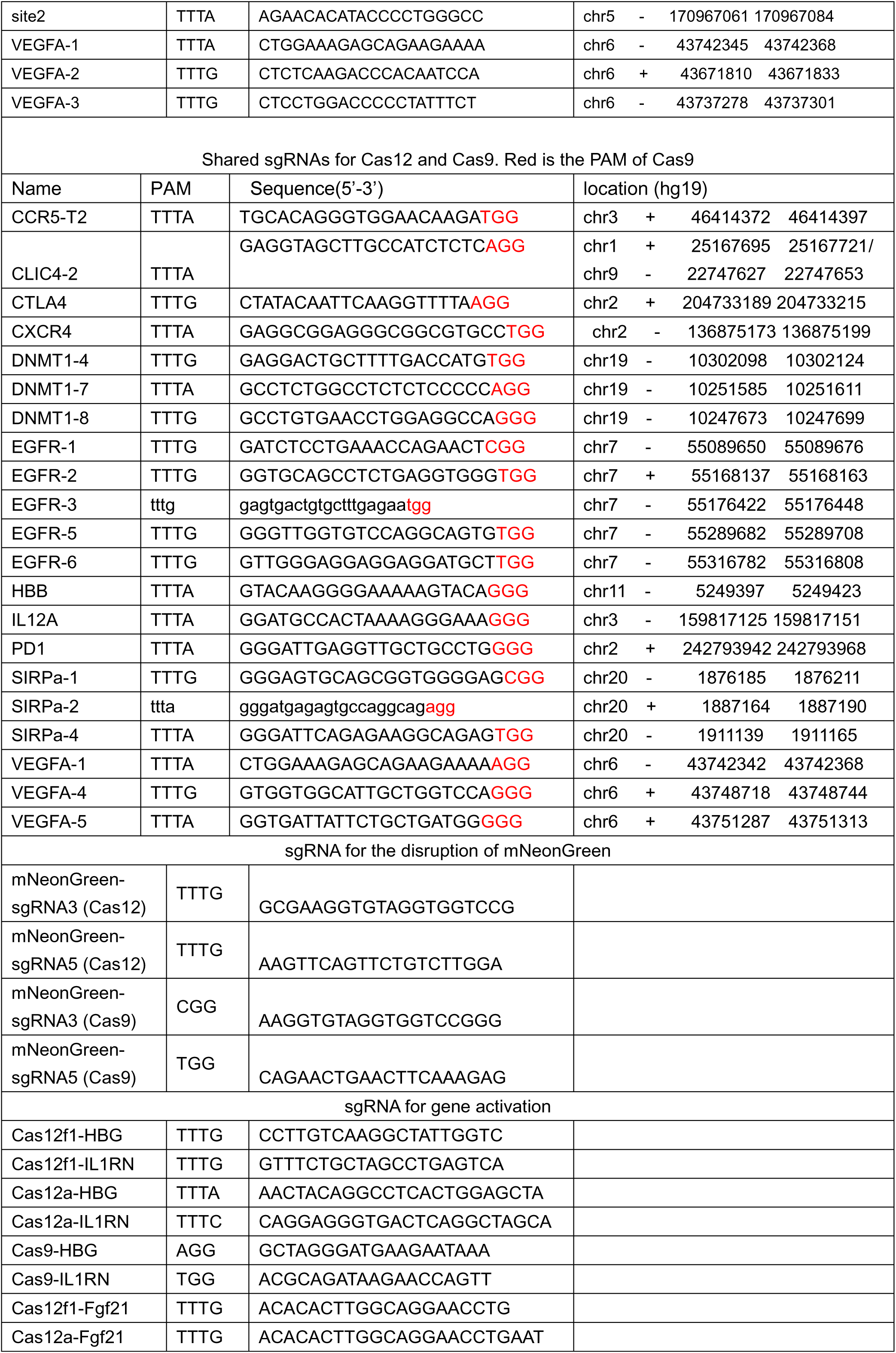
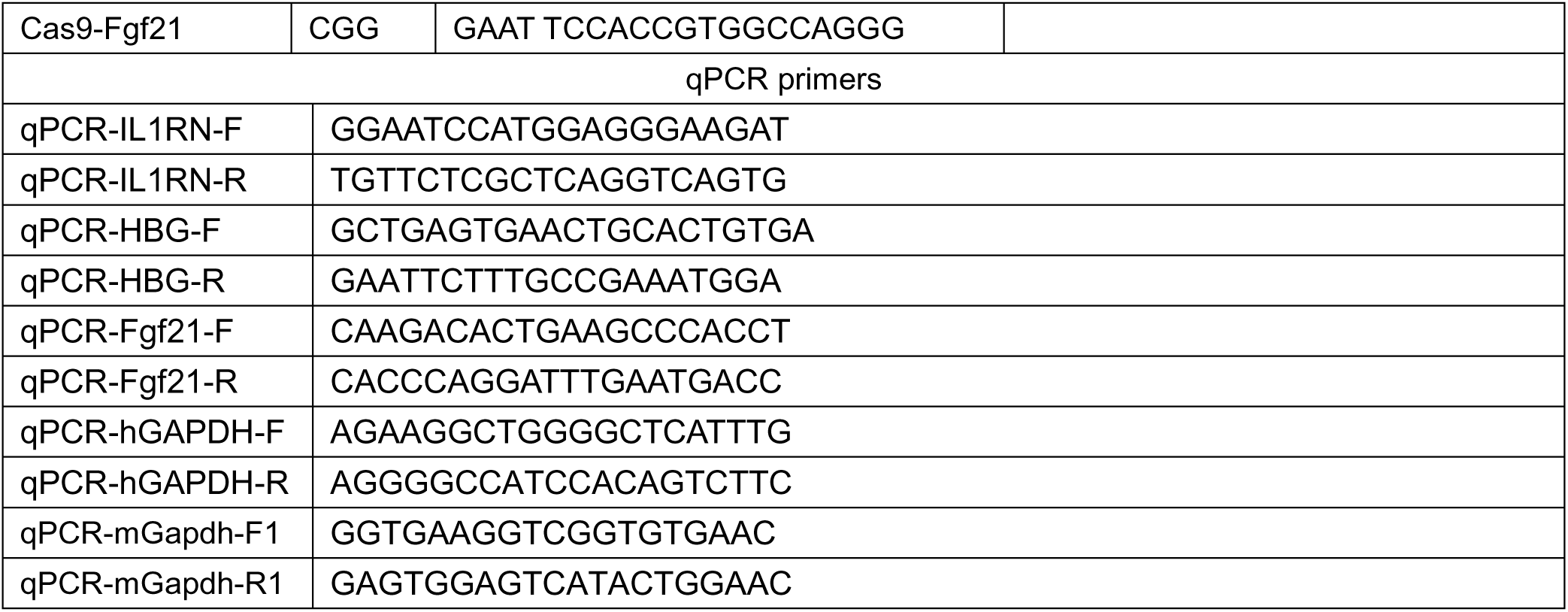
The sgRNAs and primers used in this study

**Table S3.**
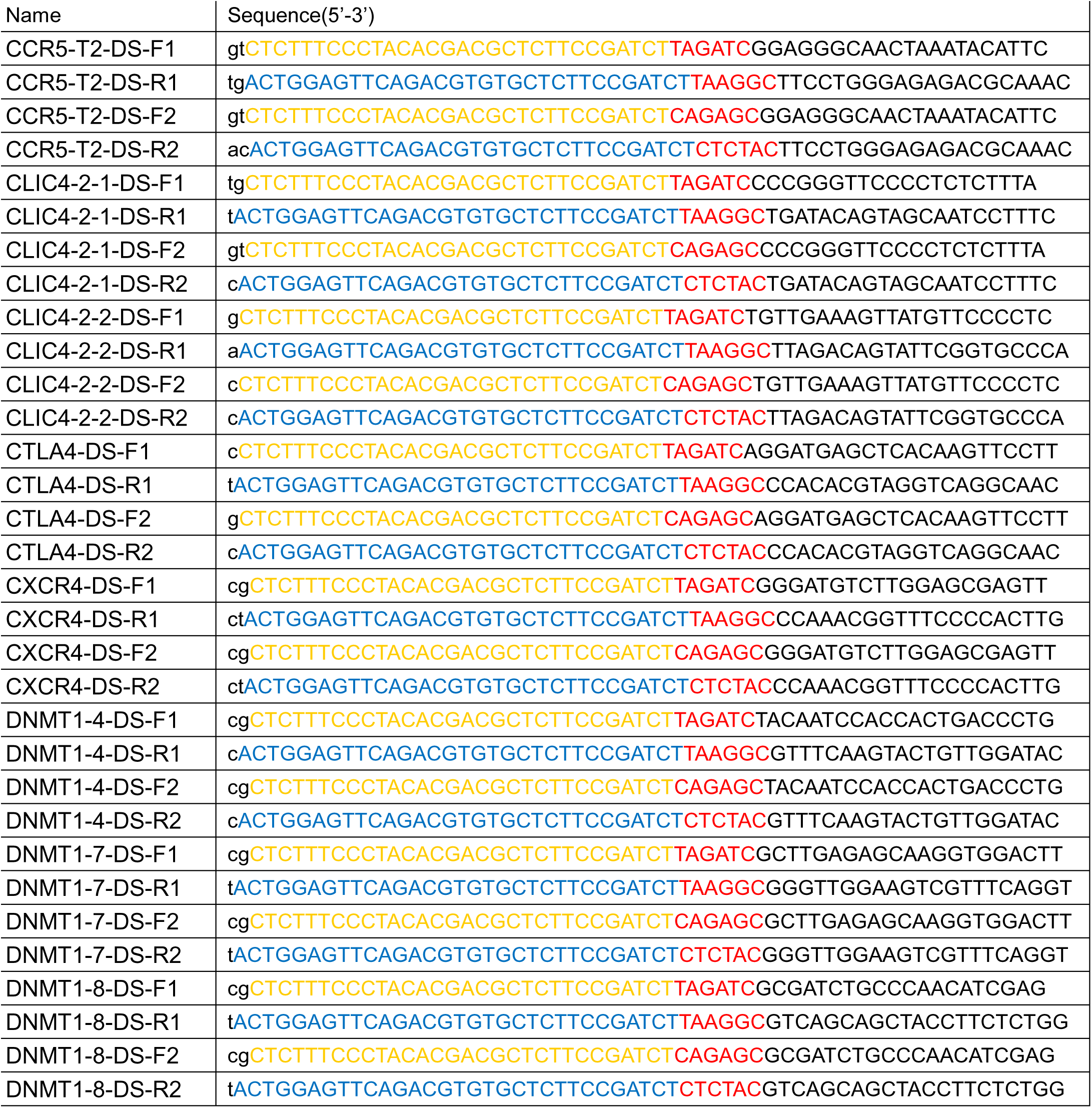
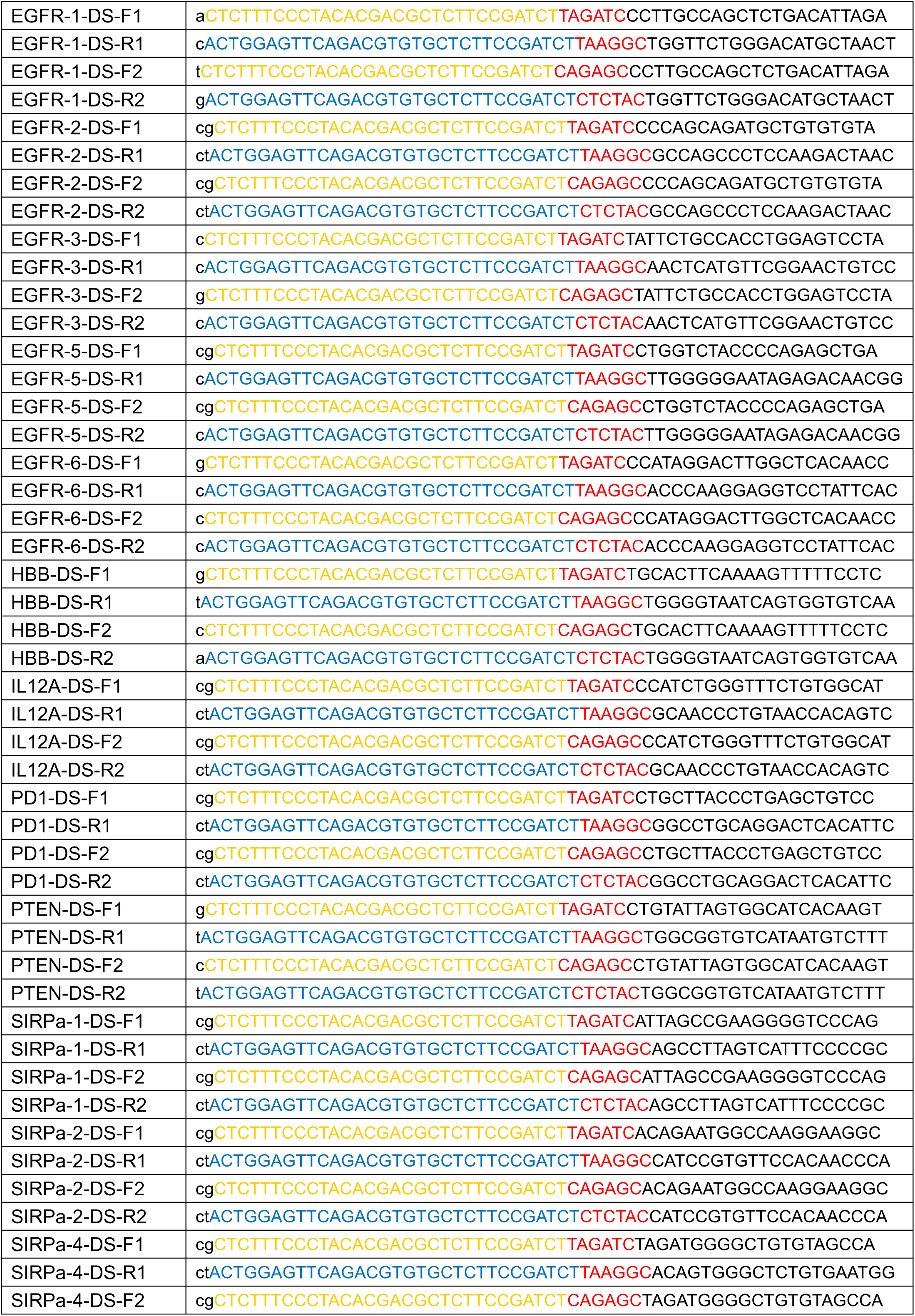
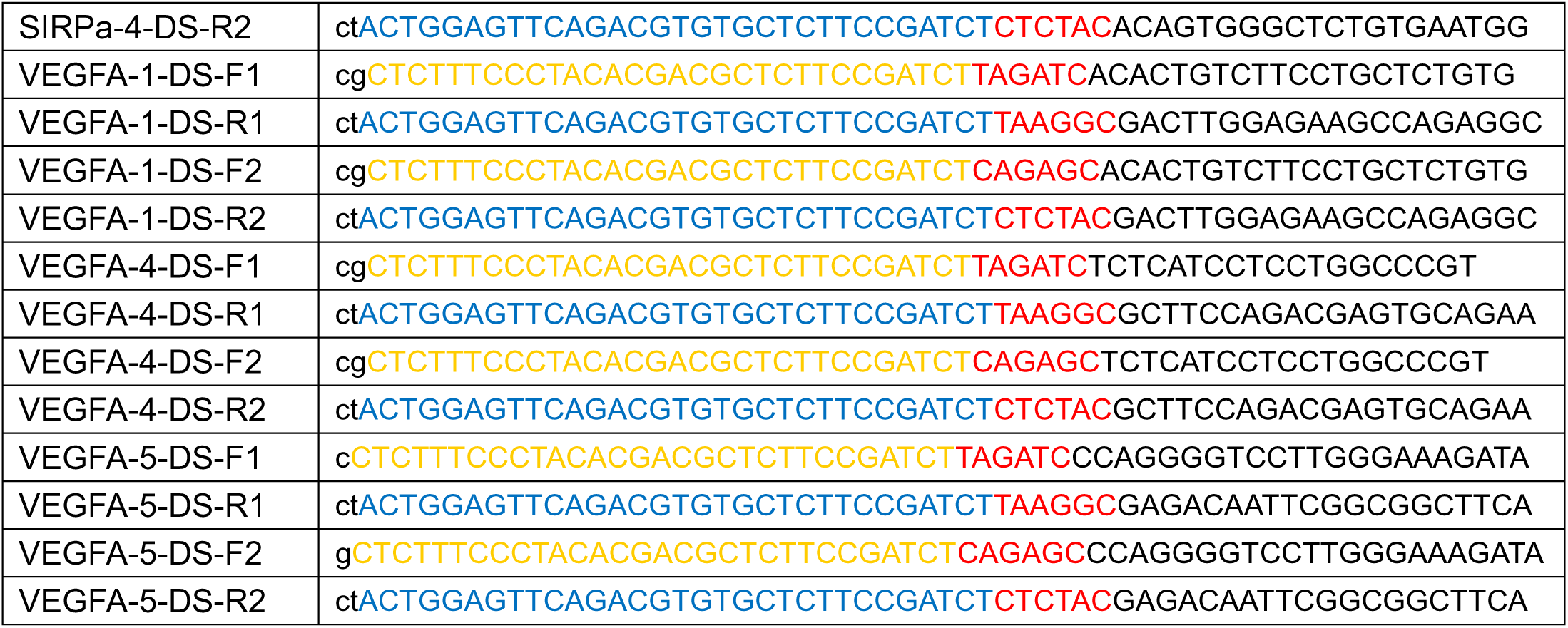
Deep-seq primers for this study

